# Control of stem cell niche and fruit development in *Arabidopsis thaliana* by *AGO10/ZWL* requires the bHLH transcription factor *INDEHISCENT*

**DOI:** 10.1101/2020.11.25.397513

**Authors:** Manoj Valluru, Karim Sorefan

**Affiliations:** Faculty of Science, University of Sheffield, Sheffield S10 2TN, United Kingdom; The Medical School, University of Sheffield, Sheffield S10 2TN, United Kingdom

**Keywords:** AGO10, ZWILLE, IND, SPT, shoot apical meristem, fruit development, auxin

## Abstract

**Background:** The shoot apical meristem (SAM) in plants is composed of a small mound of pluripotent stem cells that generate new organs. ARGONAUTE10 (AGO10) is known to be critical for maintenance of the embryonic SAM by regulating the expression of *Class III HOMEODOMAIN-LEUCINE ZIPPER* (*HD-ZIP III*) transcription factors, which then modulate downstream responses to the key phytohormone auxin. However, we do not understand how AGO10 modulates auxin responses after embryogenesis in the mature plant.

**Results:** Here we show that *AGO10* regulates auxin responses in the post-embryonic SAM via the bHLH transcription factor *INDEHISCENT* (*IND*). IND directly regulates auxin responses in the SAM regulating the auxin transporter PIN1 via direct transcriptional regulation of *PINOID* kinase. We show that a loss of function *ind* mutation significantly restores *ago10^zwl-3^* mutant SAM and fruit phenotypes. *ago10^zwl-3^* mutants overexpress *IND* and overexpression of IND phenocopies the *ago10^zwl-3^* SAM phenotypes, and regulates auxin transport and responses in the SAM. AGO10 also regulates post-embryonic development in the fruit via a similar genetic pathway.

**Conclusions:** We characterise a molecular mechanism that is conserved during post embryonic development linking *AGO10* directly to auxin responses.

## Background

All above ground tissues of angiosperm plants derive from the shoot apical meristem (SAM) and most of the global food supply is derived from these tissues, including cereals, beans and fruit. The SAM is composed of a stem cell niche that maintains a small mound of pluripotent cells and differentiated primordia that develop at the flanks of the stem cell niche [1, 2]. The SAM develops during embryogenesis and is regulated by several pathways [3] including by a balance between WUS and CLAVATA1/2/3 activity [4]. The NAC transcription factors CUP-SHAPED COTYLEDONS 1/2/3 (CUC1/2/3) are also required for meristem maintenance during and after embryogenesis to prevent organ fusions [5–7]. However, much less is known about how the SAM is maintained post embryogenesis after germination and during growth and development of the mature plant.

*ARGONAUTE10* (*AGO10*. a.k.a *ZWILLE* and *PINHEAD* and referred to as *AGO10* in this report) regulates SAM development by maintaining the stem cell niche and preventing terminal differentiation [8–10]. *ago10* mutants in the L*er* background are pleiotropic and cause SAM defects ranging from seedlings with wild-type appearance, terminally differentiated meristems with filamentous radial organs or a single radialised leaf, or empty apexes [9, 11]. Currently, we do not understand how *ago10* mutants develop such a range of phenotypes.

In contrast to the SAM phenotypes, the fruit phenotypes are fully penetrant [9, 12]. Arabidopsis develop a pod like fruit called the silique and the external tissues include the valves, replum and valve margins. The valves surround and protect the seeds and are connected to the rest of the fruit by the replum tissue. Seed dispersal is controlled by the valve margin tissues that develop next to the valves and are separated by the replum tissues. In *ago10* mutants the fruit are short, have multiple carpels and internal morphological defects [9] but we do not currently understand how *AGO10* regulates fruit development.

AGO10 maintains embryonic SAM development by regulating *HD-ZIP III* expression via an elegant small RNA mechanism [13, 14]. *HD-ZIP III* transcripts are targeted by miR165/166 and AGO10 maintains *HD-ZIP III* transcript levels by sequestering miR165/166 [13, 15, 16]. HD-ZIP III transcription factors are required for adaxial leaf domain specification and loss of function mutants cause abaxialised radial tissues to develop [17–19]. Conversely, dominant mutations in the miR165/166 target site of *HD-ZIP IIIs* cause adaxialised radial tissues to develop [20, 21]. In addition to this role in modulating HD-ZIP III gene expression, during embryogenesis *AGO10* is also required to reduce auxin signalling and probably auxin levels and this indirectly involves the auxin response transcription factor *ARF2* [22].

The radialised tissues of *ago10* mutants can also be phenocopied by overexpression of the bHLH transcription factor INDEHISCENT (IND), suggesting the *ago10* mutant phenotypes could be caused by IND overexpression. Radialisation of the SAM tissues by IND overexpression is dependent on another bHLH gene, *SPT* [23]. IND and SPT can dimerise to regulate downstream gene expression and IND directly regulates *SPT* expression [24–26]. IND is required for valve margin development and the switch from bilateral to radial symmetry during gynaecium (organ development before fertilisation) development [23, 24, 26, 27]. IND is also important for directly regulating auxin responses in this tissue. During valve margin development IND generates an auxin minima by regulating auxin transport through direct regulation of the auxin transport regulator PINOID kinase (PID) [26]. IND and AUXIN RESPONSE FACTOR3 (ARF3/ETTIN) bind auxin to regulate downstream responses [28]. The function of IND has mostly been characterised during fruit development and it is not known whether IND has endogenous functions during SAM development.

In this study, we investigated whether IND plays a role in post-embryonic SAM development and, in particular, its relationship to AGO10. We identify a signal transduction pathway that links AGO10 to auxin responses via IND, providing a new insight into the regulation of the stem cell niche in plants.

## Results

### IND mediates the *ago10^zwl-3^* SAM and fruit phenotypes

Compared to the embryonic phenotypes the characterisation of the post-embryogenic phenotypes of *ago10* have received less attention. Therefore, we characterised the early post-embryonic SAM phenotypes to obtain a better understanding of morphological changes in *ago10^zwl-3^* mutants. To investigate the effect of loss of *ago10^zwl^*^-3^ on early postembryonic SAM development we quantified the frequency of different SAM morphologies at 2 days post germination (DPG) and imaged SAM development at 3 DPG. In wild-type seedlings, the central meristem was flanked by two leaf primordia (Fig. 1A). In *ago10^zwl-3^* mutants around 32% of seedlings develop a SAM with similar morphology to wild-type seedlings (Fig. 1B, K). Around 11% of the seedlings had a single broad leaf primordia with developing trichomes and no observable morphological meristem, appearing as if two leaf primordia had fused (FP, Fig. 1C, K). Around 50% of the seedlings had one narrow primordia and no observable meristem (SP, Fig. 1D, K) and around 7% of seedlings had no observable meristem or primordia (NM, Fig. 1E, K). At 14 DPG wild-type plants had developed 4 leaves (Fig. 1F). After 14 days growth, mutations in *ago10^zwl-3^* caused a range of post-embryonic phenotypes including plants with a wild-type-like SAM (WT, Fig. 1G), cup-shaped or single leaf (CUP, Fig. 1H), pin-shaped or filamentous-like (PIN, Fig. 1I) or empty apex (EA, Fig 1J). The frequency of wild-type SAM, fused primordia, single primordia and no meristem phenotypes observed at 2 DPG correlated with the WT, CUP, PIN and EA phenotypes observed at 14 DPG respectively (Pearson r = 0.9, P value = <0.01) (Fig. 1K). This suggests that the 2 DPG phenotypes are a precursor to the 14 DPG phenotypes. Our data support the hypothesis that *AGO10* is required to maintain the stem cell niche and prevent fusion of the leaf primordia. This conclusion is supported by the observation that first two true leaves were occasionally (2/91) fused and no meristem was observed (Additional file 1: Figure S1).

**Fig. 1.**
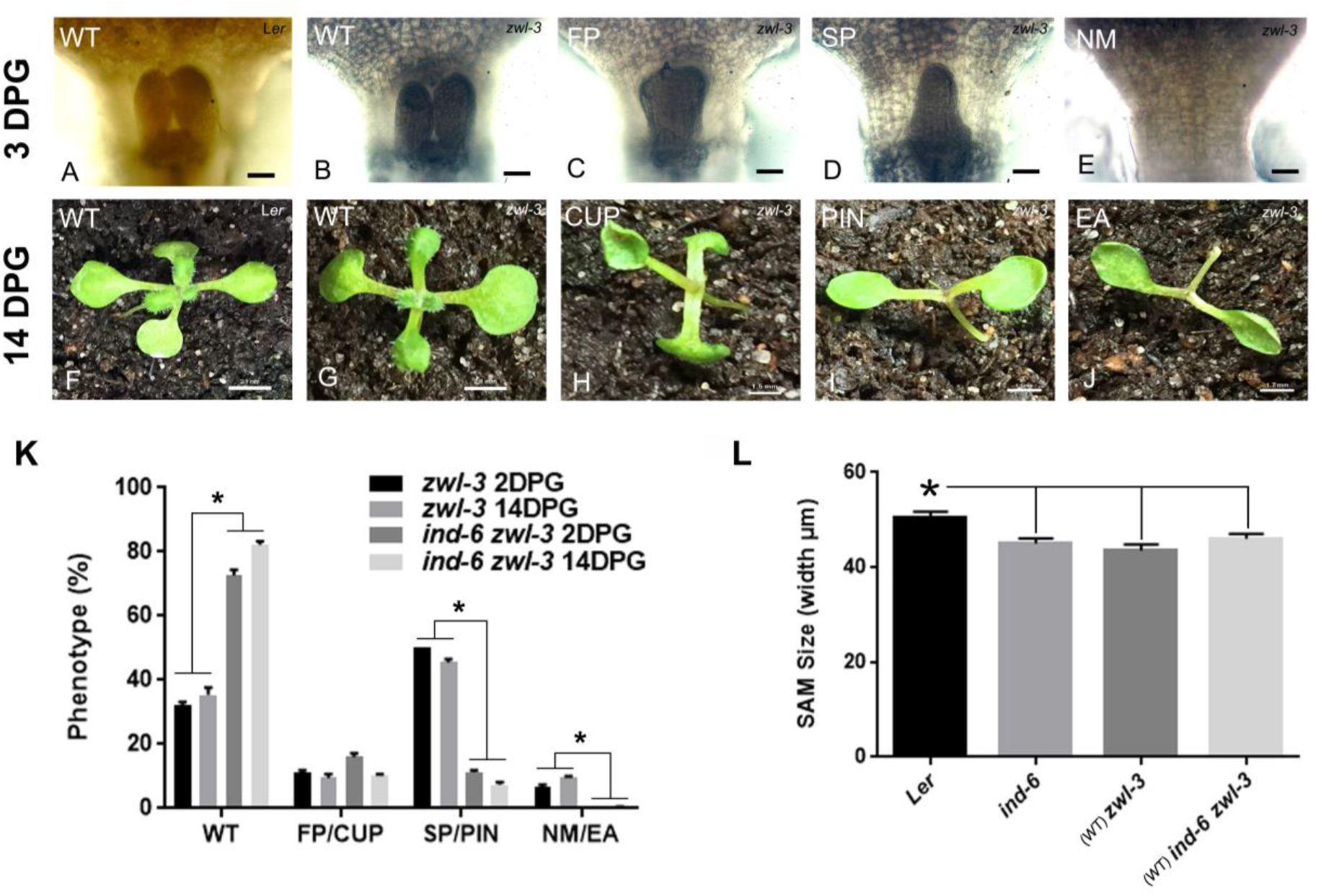
Post embryonic meristem defects in *ago10^zwl-3^* are restored by the *ind-6* mutation. (A) Wild type SAM at 3 DPG. (B-E) SAM development of *ago10^zwl-3^* seedlings grown for 3 DPG develop either a wild-type like SAM (WT), fused primordia (FP), single primordia (SP) or no observable meristem (NM). (F) Wild type SAM at 14 DPG. (G-J) SAM development of *ago10^zwl-3^* seedlings grown for 14 DPG develop either a wild-type like SAM (WT), single leaf or cup-shaped leaf (CUP), pin-shaped or filamentous-like (PIN) or an empty apex (EA). Scale bar for 3 DAG = 50 μm. (K) The frequency of SAM phenotypes in *ago10^zwl-3^, ago10^zwl-3^,ind-6* double mutants after 3 and 14 DPG. The frequency of WT, FP, SP and NM phenotypes at 2 DPG and WT, CUP, PIN and EA phenotypes at 14DPG respectively is closely correlated. The frequency of PIN and EA phenotypes in *ago10^zwl-3^* is reduced in the *ago10^zwl-3^,ind-6* double mutant (n=3 biological replicates). Values are means ± SE. Tukey’s multiple comparisons test (*ind-6 zwl-3* vs. *zwl-3*), *p<0.001. (L) Width of SAM in seedlings with WT phenotype after 3 DPG in wild type, *ind-6, ago10^zwl-3^* and *ago10^zwl-3^,ind-6* double mutants (n=10). Values are means ± SE. Tukey’s multiple comparisons test, *p<0.05 (L*er* vs. mutants).

We then asked whether the *ago10^zwl-3^* phenotypes were dependent on *IND* function by scoring the different phenotypes of *ago10^zwl-3^, ind-6* and double mutants after 14 DPG. *ind-6* is a L*er* ecotype enhancer trap line and is considered to be a null allele [29]. 100% of wild-type and *ind-6* mutant seedlings had a WT SAM phenotype suggesting IND does not severely effect SAM development (n=50 and 3 biological replicates respectively). A key finding was that the *ind-6* mutation partially restored the *ago10^zwl-3^* mutant phenotypes (Fig 1K), demonstrating the *ago10^zwl-3^* phenotypes are dependent on *IND* function. In the double mutant the frequency of the *ago10^zwl-3^* PIN and EA SAM defects were significantly reduced compared to the *ago10^zwl-3^* single mutants, however the frequency of CUP phenotypes was not significantly changed (Fig. 1K). Our data suggests that the *ago10^zwl-3^* PIN and CUP phenotypes are dependent on *IND* function.

Although the *ind-6* mutant did not have obvious SAM defects, a microscopic analysis of *ind-6* mutants with WT phenotypes showed that the meristem size was significantly reduced compared to wild type (Fig. 1L). This suggests that *IND* may be required to maintain the size of the stem cell niche. The meristem size of double mutants with the WT phenotype was not significantly different from *ind-6* or *ago10^zwl-3^* mutants, suggesting loss of *IND* does not does rescue the *ago10^zwl-3^* mutant phenotype by affecting meristem size (Fig. 1L).

To further support our hypothesis that *ago10^zwl-3^* phenotypes are dependent on *IND*, we analysed whether the *ind* mutation could rescue the *ago10^zwl-3^* phenotypes in another developmental context. In comparison to the SAM phenotypes, fruit development is particularly sensitive to loss of *AGO10* as all *ago10^zwl-3^* fruit are short [9, 12]. However, the molecular mechanism causing these fruit phenotypes is not known. Since *IND* regulates fruit development, we analysed whether the *ago10^zwl-3^* mutant fruit phenotypes were mediated by *IND*. *IND* mutants develop significantly longer fruit than wild-type plants [30], and loss of *IND* function increases replum width [31]. In agreement with previous findings, we found *ind-6* mutants have significantly longer fruit (Fig. 2A, B) and increased replum width (Fig. 2C) compared to wild-type plants (Fig. 2A-C). *ago10^zwl-3^* mutants have short fruit (Fig. 2A, B) and reduced replum width (Fig. 2C), and loss of *IND* function in this background partially restored these phenotypes (Fig. 2A-C). One interpretation of the data is that *AGO10* represses *IND* to promote separation/prevent fusion of the valve margins.

**Fig. 2.**
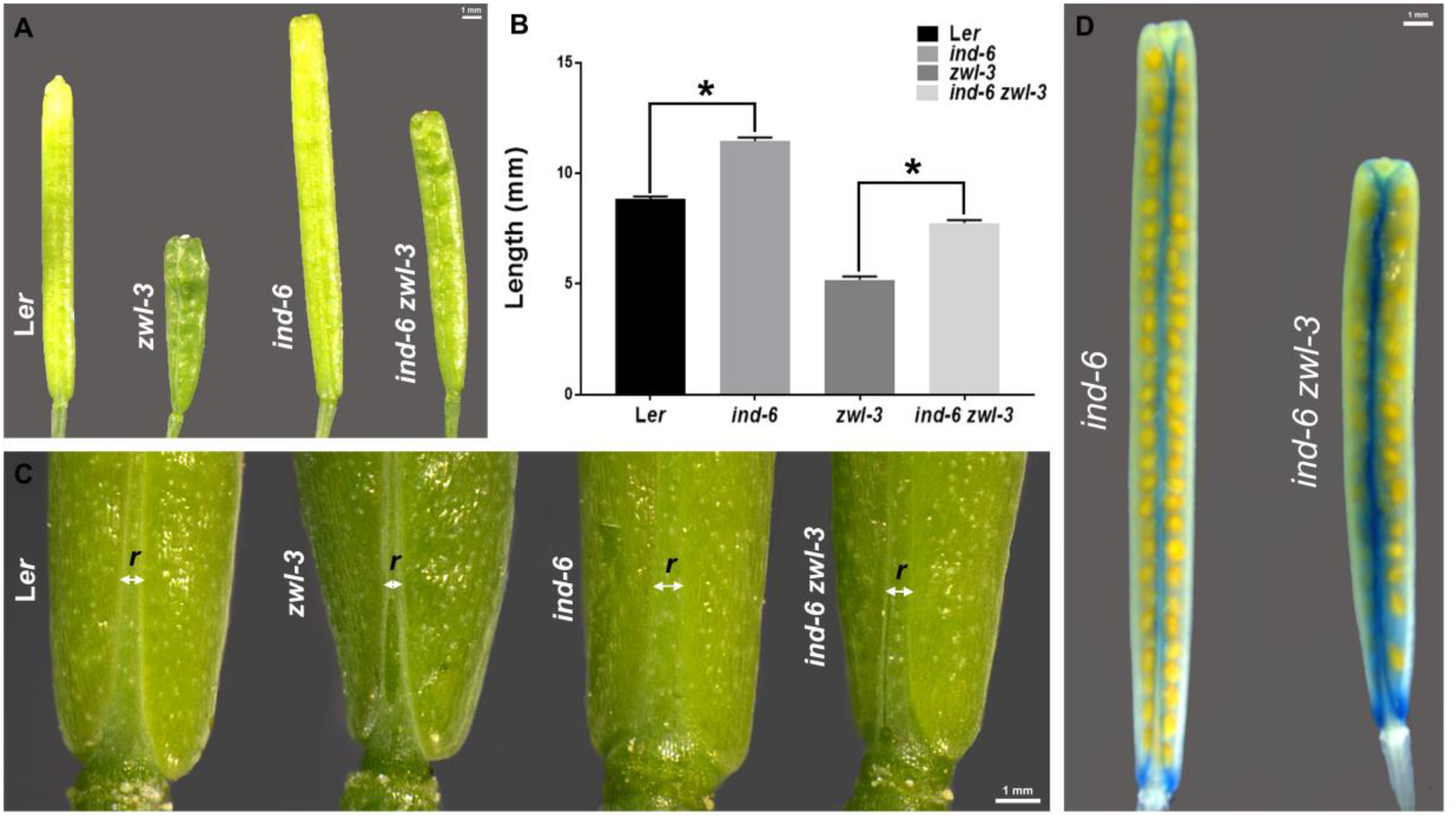
The fruit development defects in *ago10^zwl-3^* are restored by the *ind-6* mutation and *IND* expression is increased *ago10^zwl-3^* as visualised by the *ind-6* GUS reporter line. (A) The length of the mature fruit (stage 17) of L*er* (wild type), *ind-6, ago10^zwl-3^* and *ago10^zwl-3^,ind-6* double mutants. (B) Quantification of mature fruit length (n=40, Values are means ± SE, Tukey’s multiple comparisons test, *p<0.05). (C) *AGO10* and *IND* regulate replum (r) width. White line with double arrows indicates repla of L*er, ind-6, ago10^zwl-3^* and *ago10^zwl-3^,ind-6* double mutants mature fruits. (D) GUS staining in mature fruit of the *ind-6* GUS reporter line is localised to the valve margins between the valves and replum. Compared to the *ind-6* single mutant, GUS expression was increased in the *ago10^zwl-3^,ind-6* double mutant.

Our data suggest the *ago10^zwl-3^* mutant phenotypes are dependent on *IND* function in both the SAM and fruit, which demonstrates that a genetic pathway involving *AGO10* and *IND* is conserved in both organs.

The smaller replum width in *ago10^zwl-3^* mutants is indicative of increased IND expression [32], so we tested whether *IND* expression was increased in *ago10^zwl-3^* mutant fruit. To investigate the expression of *IND* in the fruit of double mutants, we utilised the fact that the *ind-6* mutant carries an enhancer trap with a beta-glucuronidase (GUS) reporter [29]. GUS expression in *ind-6* mutants faithfully reported *IND* expression in the fruit [29]. We observed that GUS expression was increased in the fruit of double mutants compare to *ind-6* single mutants (Fig. 2D). This provided the first evidence that *AGO10* is required to repress *IND*, and we hypothesised a similar mechanism exists during SAM development.

### AGO10 regulates proper *IND* and *SPT* expression during SAM development

We used *pIND::GUS, pIND::IND-YFP, pAGO10:: YFP-AGO10* lines and quantitative reverse transcriptase PCR (qRT-PCR) to characterise *IND* expression in seedlings. First, we investigated whether AGO10 and IND were expressed in the same tissues during seedling and fruit development. At 3 DPG AGO10 was observed in the meristem and adaxial domain of leaf primordia of *pAGO10:: YFP-AGO10* lines (Fig. 3A), in agreement with a previous finding [33]. Histochemical staining of *pIND::GUS* lines suggest IND was weakly expressed in the meristem and leaf primordia (Fig. 3C, D), and very weak IND-YFP expression also observed in the meristem and leaf primordia (Fig. 3B, Additional file 1: Figure S2). IND was also found to be expressed in the vegetative meristem as measured by Tiling Array Express data (Additional file 1: Figure S2) [34].

**Fig. 3.**
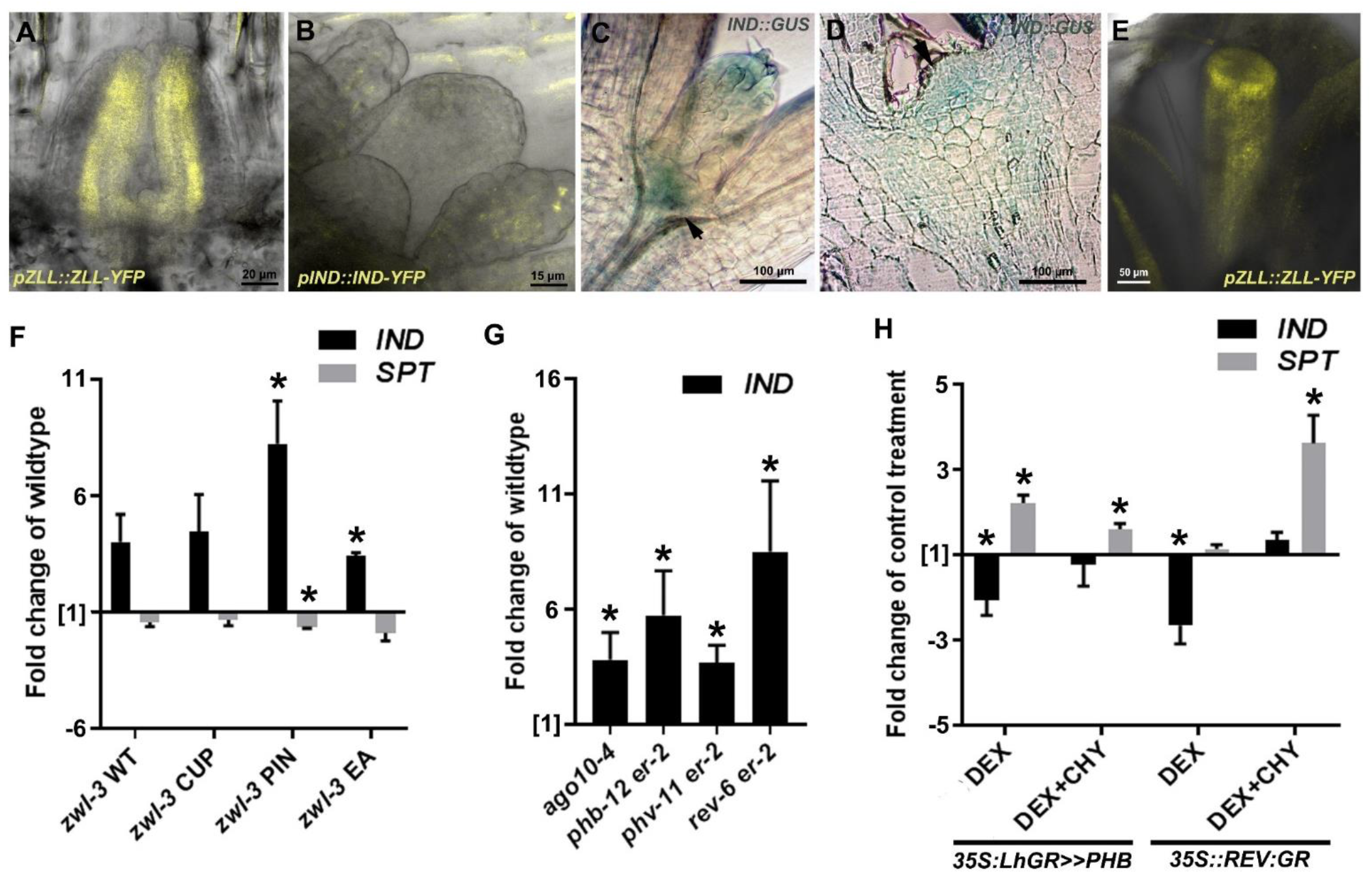
*AGO10*, *IND* and *SPT* expression during post-embryonic development. (A) Expression of *pAGO10::YFP-AGO10* in the SAM at 3DPG. (B) Expression of *pIND::IND-YFP* in the SAM at 7 DPG. (C) Whole mount GUS staining of *pIND::GUS* reporter line in the SAM at 3 DPG. (D) Cross section through SAM of *pIND::GUS* line stained for GUS. Arrows indicate meristem region. (E) Expression of *pAGO10::YFP-AGO10* in stage 9 gynoecium showing YFP signal in the presumptive valves, valve margins and replum. (F) qRT-PCR analysis of *IND* and *SPT* expression in each of the phenotypic groups of *ago10^zwl-3^* mutants. *IND* expression is significantly increased in PIN and EA phenotypes of *ago10^zwl-3^* mutants and SPT was significantly repressed in seedlings with PIN phenotypes (fold change of wild-type. n=2). (G) qRT-PCR analysis of *IND* expression in 7-day-old *ago10-4, phb-12 er-2, phv-11 er-2* and *rev-6 er-2* seedlings (n=3). *IND* expression is significantly increased in all mutants. (H) qRT–PCR in 7-day-old *35S::LhGR>>PHB* and *35S::REV:GR* seedlings ±DEX and ±cycloheximide (CHY) (n=3). PHB and REV indirectly downregulates *IND* expression and directly upregulates *SPT* expression. *p<0.05 two tailed t test. Values are means ± SE.

During fruit development IND has been shown to be expressed as early as stage 9 [24, 26, 28] and we observed AGO10-YFP expression in the presumptive valves, replum and style of stage 9 gynoecia (Fig. 3E). We did not observe *pIND::GUS* or *pIND::IND-YFP* expression during embryo development, which suggests IND does not function during embryogenesis or that the expression of the IND reporters were not sensitive enough to detect expression in the embryo. The expression data suggests that AGO10 and IND are expressed in overlapping tissue domains during post-embryonic SAM and gynoecium development.

We next asked whether *AGO10* and *HD-ZIP IIIs* regulate *IND* expression using qRT-PCR. *IND* expression was increased in *ago10^zwl-3^* mutants in the L*er* background (Fig. 3F) and to a lesser extent in the *ago10-4* mutant in the Col-0 background (Additional file 1: Figure S3). Our data suggests *AGO10* is a negative regulator of *IND* in both Col and L*er* backgrounds, but in the *ago10-4* mutant (Col background) IND expression did not reach a high enough threshold to cause SAM defects or alternatively other factors suppress the effects of IND overexpression. We quantified *IND* expression in the different *ago10^zwl-3^* phenotypes to investigate whether there was a correlation between expression level and phenotypes. We found *IND* expression was significantly increased in *ago10^zwl-3^* mutants with PIN and EA phenotypes, which were the phenotypes that were rescued in the *ago10^zwl-3^,ind-6* double mutant (Fig. 3F and 2C). This data suggest that the PIN and EA phenotypes but not the CUP phenotype were correlated with increased IND expression (Fig. 1C).

How does AGO10 regulate *IND* expression? Since AGO10 binds small RNAs one possibility is that AGO10 regulates *IND* via a microRNA dependent mechanism. However, this is probably not the case because we did not identify any microRNAs in miRBase that would be predicted to target *IND*, and *IND* expression was not strongly changed in microRNA biogenesis mutants (Additional file 1: Figure S4). Another possibility is that HD-ZIP III’s may regulate *IND*. To test this hypothesis, cDNA was prepared from 14-day old wild-type, *ago10-4, phb-12, phv-11* and *rev-6* seedlings and gene expression was quantified using qRT-PCR. When compared to wild-type, there was a significant increase of *IND* expression in *phb-12, phv-11* and *rev-6* (Fig. 3G). Upregulation of *IND* in the *hd-zip III* mutants suggests that *PHB*, *PHV*, and *REV* negatively regulate *IND*. Consistent with this hypothesis, *IND* was repressed by DEX induction of PHB and REV in the *35S:LhGR>>PHB and 35S::REV-GR* transgenic lines respectively (Fig. 3H) [35, 36]. However, in the presence of cycloheximide (CHY), which inhibits protein synthesis, PHB and REV induction did not induce *IND* (Fig. 3H), suggesting the PHB and REV regulate *IND* indirectly.

*IND* function is dependent on *SPATULA* (*SPT*) [23] [24–26], so we then investigated whether AGO10 may also regulate SPT expression via PHB and REV. *SPT* expression was induced by PHB and REV induction in the *35S:LhGR>>PHB and 35S::REV:GR* transgenic lines when protein synthesis was blocked, suggesting PHB and REV directly regulated *SPT* (Fig. 3H). In addition, analysis of REV ChIP-seq datasets also identified *SPT* as a potential target of REV (GSE26722, Additional file 1: Figure S5) [37]. Although we did not test whether PHV can bind *SPT* gene in this study, global binding studies using DAP-seq show recombinant PHV protein can bind the *SPT* promoter (GSM1925338, Additional file 1: Figure S5) [38]. Consistent with this, we found the levels of *SPT* expression was reduced in *ago10^zwl-3^* mutant seedlings (Fig. 3F), where we show HD-ZIP III (PHB, PHV and REV) expression was reduced (Additional file 1: Figure S3).

Taken together, our data suggests *AGO10* and *IND* are expressed in the same tissues in the SAM, and that *AGO10* and *HD-ZIP IIIs* are required to repress *IND*. AGO10 may also upregulate *SPT* gene expression via direct regulation by HD-ZIP IIIs, which could be a potential mechanism to modulate IND activity.

### IND functions in a network with AGO10 and HD-ZIP III

We also investigated whether IND may also regulate the expression of other components of the AGO10 pathway including *AGO10*, miR165/166 or *HD-ZIP IIIs* (*PHB*, *PHV* and *REV*).

We tested whether overexpression of IND affected *AGO10* expression and we found DEX induction of IND-GR repressed *AGO10* expression using qRT-PCR (Fig, 4A). Overexpression of IND also repressed *AGO10* when protein synthesis was blocked suggesting IND regulated AGO10 directly (Fig, 4A). Chromatin immunoprecipitation (ChIP) followed by qPCR confirmed that IND-GR or a complex including IND-GR bound the *AGO10* promoter directly (Fig, 4B).

**Fig. 4.**
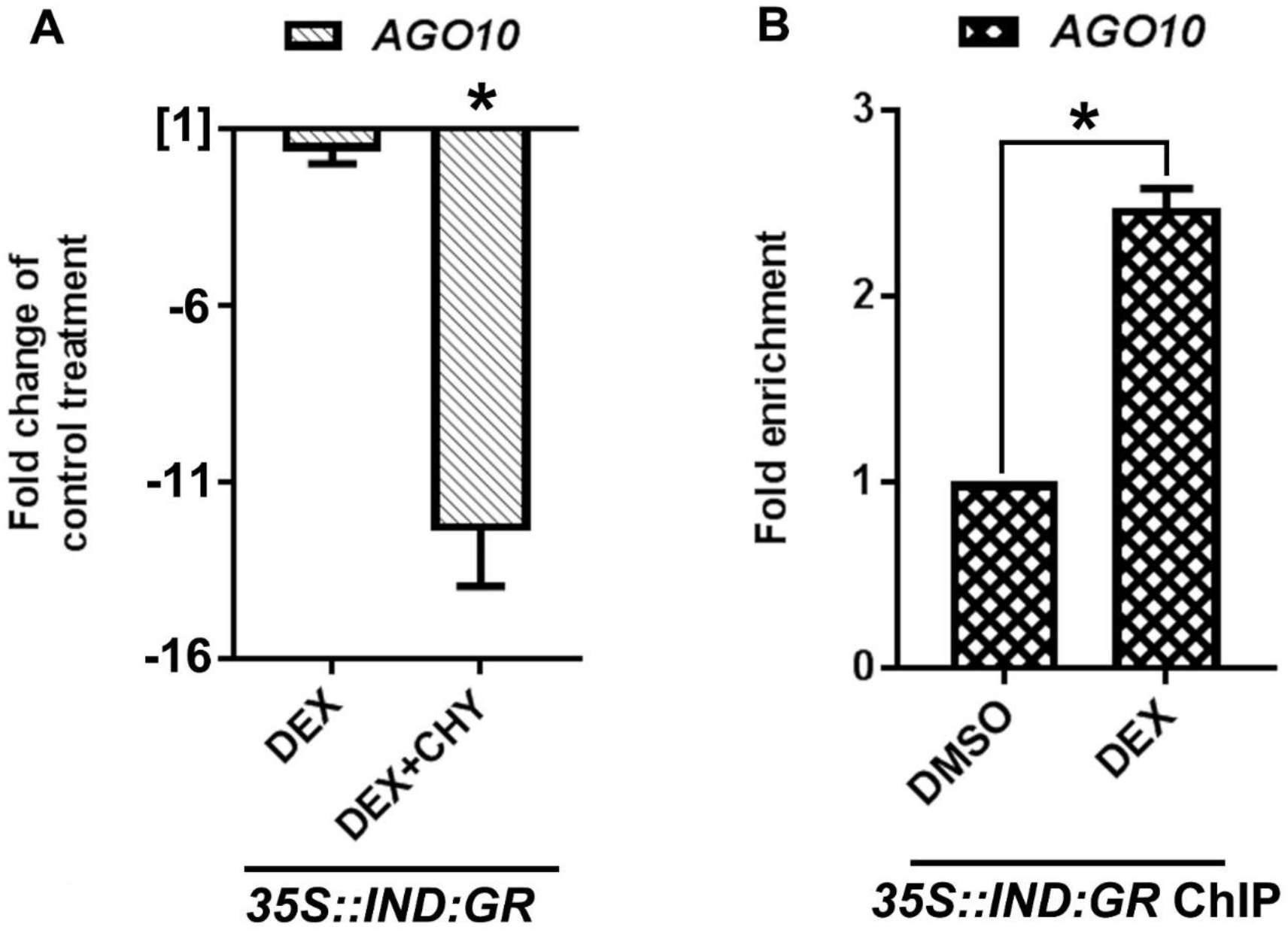
IND directly regulates *AGO10* expression. (A) qRT–PCR in 7-day-old *35S::IND-GR* seedlings ±DEX and ±cycloheximide (CHY) (n=3). IND directly downregulates *AGO10* expression (two tailed t test, *p<0.05). (B) ChIP using anti-GR antibody followed by qPCR using *35S::IND:GR* line. IND-GR binds an upstream element in the *AGO10* promoter (−926-1175 bp) that encodes a putative IND binding site (n=3, two tailed t test, *p<0.05). Values are means ± SE.

We next tested whether IND affected miR165/166 expression. sRNA-seq analysis showed that, in agreement with previous studies, the levels of miR165a-b and miR166a-g were increased in *ago10^zwl-3^* mutants (Table 1). miR165/166 levels were not significantly changed in *ind-6* single mutants compared to wild type (Table 1). In the *ago10^zwl-3^ ind-6* double mutant the expression of miR166c-g were slightly reduced compared to *ago10^zwl-3^* (Table 1). Northern blot analysis suggested miR166a levels were not significantly different between *ago10^zwl-3^* and double mutants (Additional file 1: Figure S6). Although we used a probe designed to bind miR166a it was likely to bind multiple members of this family because they have very similar sequences. Therefore, *IND* may regulate SAM development by regulating miR166 levels, but it is not known how *IND* may regulate miRNA166 levels. A possible mechanism that would require testing would be that IND regulates the transcription of the pri-miRNAs.

**Table 1:**
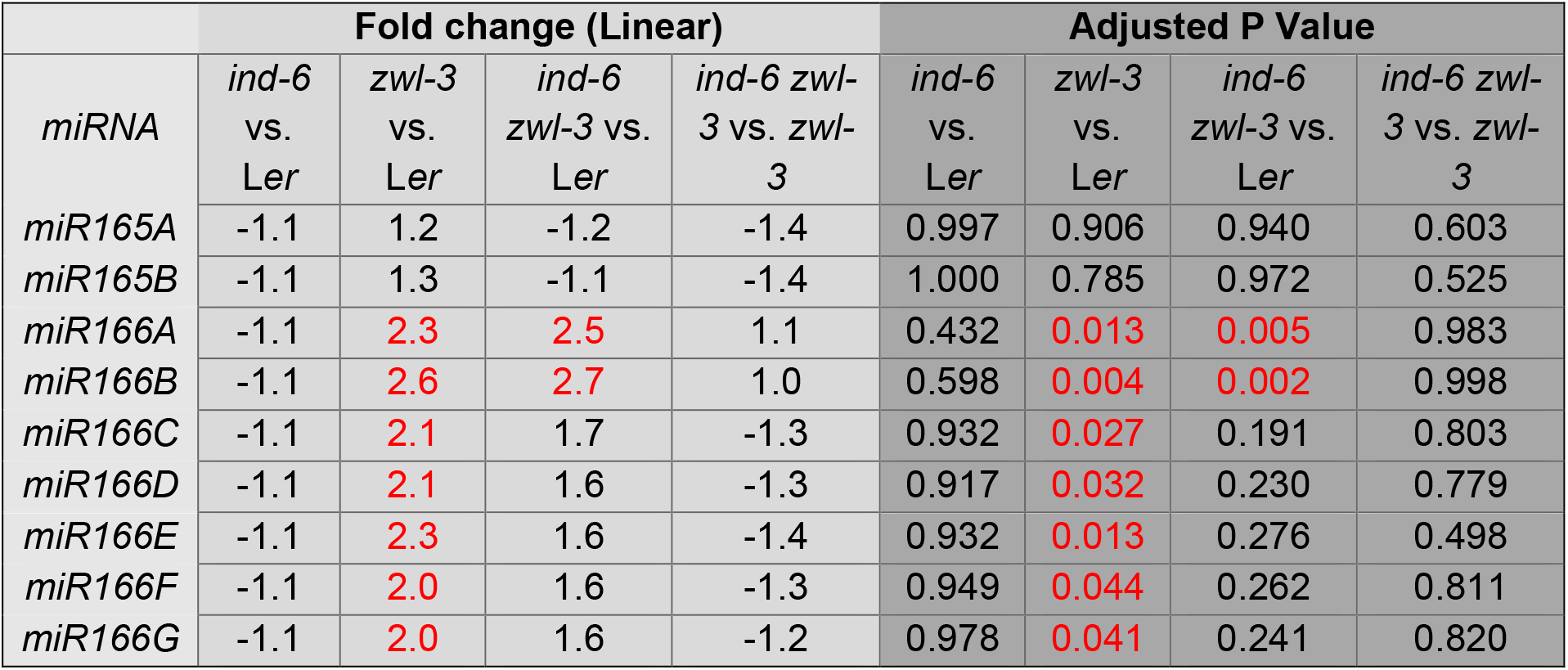
Expression of miRNA 165/166 family in *ago10^zwl-3^, ind-6* mutants *and ago10^zwl-3^ ind-6* double mutant from sRNAseq data (n=2). Fold change and corresponding adjusted P values are shown.

| miRNA | Fold change (Linear) |  |  |  | Adjusted P Value |  |  |  |
| --- | --- | --- | --- | --- | --- | --- | --- | --- |
|  | *ind-6*<br>vs.<br>Ler | *zwl-3*<br>vs.<br>Ler | *ind-6*<br>*zwl-3* vs.<br>Ler | *ind-6 zwl-3*<br>vs. *zwl-3* | *ind-6*<br>vs.<br>Ler | *zwl-3*<br>vs.<br>Ler | *ind-6*<br>*zwl-3* vs.<br>Ler | *ind-6 zwl-3*<br>vs. *zwl-3* |
| *miR165A* | -1.1 | 1.2 | -1.2 | -1.4 | 0.997 | 0.906 | 0.940 | 0.603 |
| *miR165B* | -1.1 | 1.3 | -1.1 | -1.4 | 1.000 | 0.785 | 0.972 | 0.525 |
| *miR166A* | -1.1 | 2.3 | 2.5 | 1.1 | 0.432 | 0.013 | 0.005 | 0.983 |
| *miR166B* | -1.1 | 2.6 | 2.7 | 1.0 | 0.598 | 0.004 | 0.002 | 0.998 |
| *miR166C* | -1.1 | 2.1 | 1.7 | -1.3 | 0.932 | 0.027 | 0.191 | 0.803 |
| *miR166D* | -1.1 | 2.1 | 1.6 | -1.3 | 0.917 | 0.032 | 0.230 | 0.779 |
| *miR166E* | -1.1 | 2.3 | 1.6 | -1.4 | 0.932 | 0.013 | 0.276 | 0.498 |
| *miR166F* | -1.1 | 2.0 | 1.6 | -1.3 | 0.949 | 0.044 | 0.262 | 0.811 |
| *miR166G* | -1.1 | 2.0 | 1.6 | -1.2 | 0.978 | 0.041 | 0.241 | 0.820 |

During embryogenesis it has previously been shown that the *AGO10* mutant phenotypes are due to reduced *HD-ZIPIII* expression [13]. Therefore we tested whether IND rescues the *ago10^zwl-3^* mutant phenotypes by restoring *HD-ZIP-III* expression after embryogenesis. First, we investigated the expression of *HD-ZIP III* expression in the different phenotypic groups of *ago10^zwl-3^* mutants. We identified a correlation between phenotype and *HD-ZIP III* expression levels. *PHB*, *PHV* and *REV* expression were reduced in *ago10^zwl-3^* mutants with PIN and EA phenotypes compared to wild-type plants (Additional file 1: Figure S3). Therefore, *AGO10* is required to maintain *PHB*, *PHV* and *REV* expression, and seedlings with reduced *HD-ZIP III* expression can sometimes develop a WT SAM phenotype. In the *ind-6* mutant *HD-ZIP III* expression was not significantly different from wild type (Additional file 1: Figure S3) and we found in the double mutant *PHB, PHV* and *REV* expression were not significantly different to *ago10^zwl-3^* single mutants (Additional file 1: Figure S3). This suggests that IND probably does not regulate *HD-ZIP III* expression.

In summary, *IND* may function in a feedback loop to repress *AGO10* and possibly upregulate some members of the miR166 family.

### Overexpression of IND disrupts auxin responses and meristem associated gene expression

We have shown that AGO10 regulated IND expression to maintain stem cell niche development, but we do not understand which processes were regulated downstream of IND. An obvious candidate would be that AGO10 and IND regulate auxin responses because both genes have been linked to the regulation of auxin signalling and transport. During embryogenesis AGO10 is required to maintain proper auxin responses, but the molecular mechanism is not fully understood. During fruit development, IND regulates auxin transport in the valve margin tissues by repressing *PID* and inducing *WAG2* expression at the valve margins, which leads to PIN relocation from apico-basal to apolar-lateral [26]. Therefore, we investigated whether overexpression of IND regulates auxin responses in the SAM as it does in the fruit.

To investigate the effect of IND overexpression on auxin transport we observed the localisation of the auxin transporter, *pPIN1::PIN1-GFP*, after DEX induction in the IND-GR lines using confocal microscopy. In mock-treated seedlings PIN1-GFP expression was polarly localised in the older leaf primordia. In younger leaf primordia PIN1-GFP localisation was more diffuse, and expression was restricted to the proximal end of the primordia (Fig. 5C). After DEX induction of IND-GR (18h), the PIN1-GFP signal was reduced and polar localisation was lost (Fig. 5D). As we have shown previously, IND-GR induction did not affect *PIN1* gene expression but did directly repress *PID* (Additional file 1: Figure S7) [26]. Therefore, we suggest that in the SAM, as in during gynoecium development, ectopic expression of IND affects the levels and polar localisation of PIN1 through transcriptional regulation of *PID*.

**Fig. 5.**
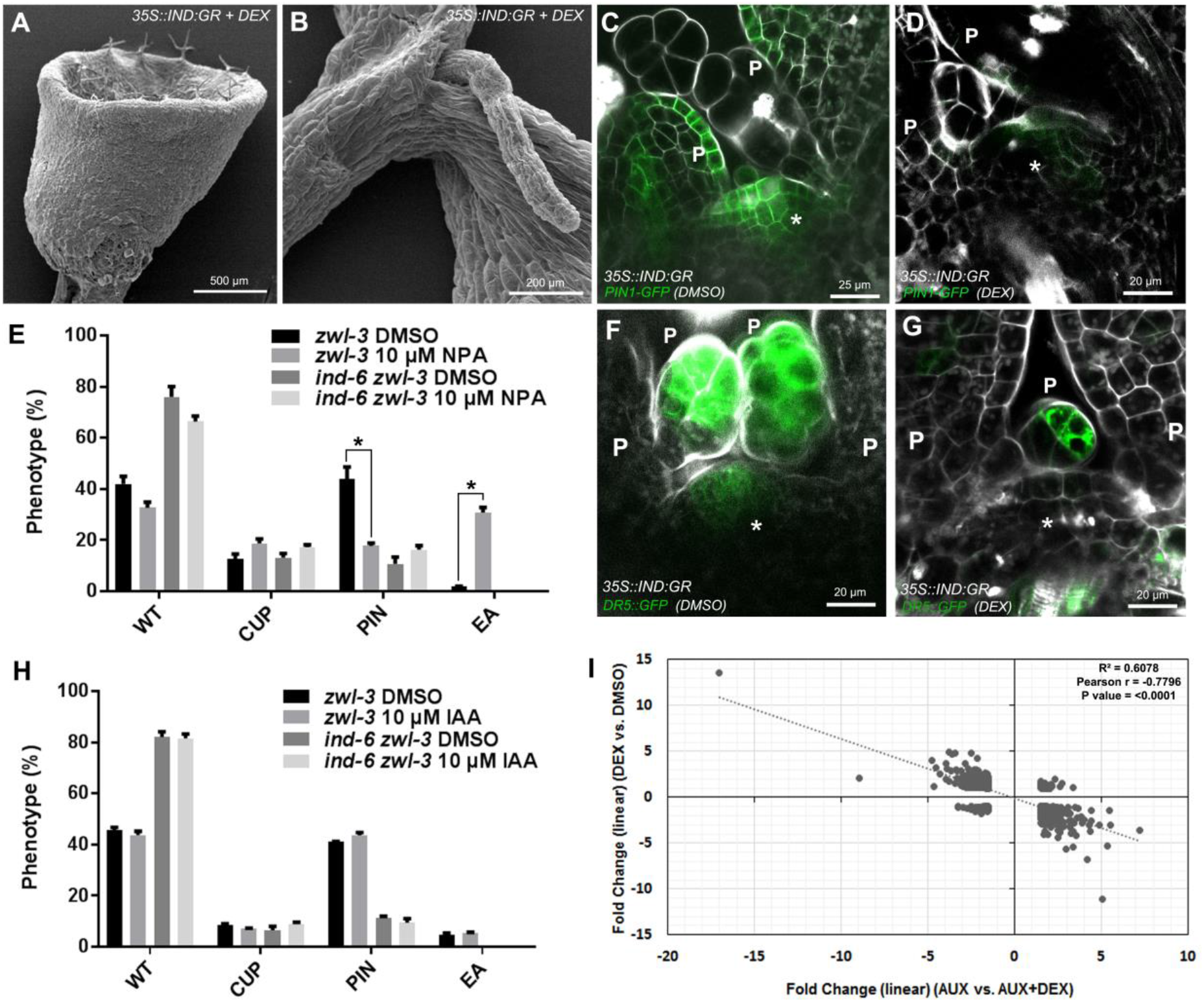
Overexpression of IND phenocopies *ago10^zwl-3^* and alters auxin responses. *35S::IND:GR* seedlings germinated on 10μm DEX for 21 days develop (A) CUP and (B) PIN phenotypes. (C) PIN1-GFP is polarly localised in leaf primordia (p) of *35S::IND:GR,pPIN1::PIN1:GFP* lines after 18h mock treatment (DMSO). (D) PIN1-GFP levels are reduced and polar localisation is lost after IND-GR induction in *35S::IND:GR,pPIN1::PIN1:GFP* lines by 18h 10μM DEX treatment. (E) Inhibiting auxin transport of *ago10^zwl-3^* seedlings after 14 DPG treatment with NPA reduces the frequency of PIN and increases the frequency EA phenotypes. NPA treatment does not significantly affect the frequency of mutant phenotypes in *ago10^zwl-3^,ind-6* double mutants (n=50, 3 biological replicates, Tukey’s multiple comparisons test, *p<0.05). (F) Expression of the auxin signalling reporter *DR5rev::GFP* in mock treated *35S::IND:GR,DR5rev::GFP* seedlings and (G) after 18h 10μM DEX treatment. (H) Exogenous auxin treatment for 21 days 10μM does not affect the frequency of SAM phenotypes in *ago10^zwl-3^* or *ago10^zwl-3^,ind-6* double mutants (n=50, 3 biological replicates, Tukey’s multiple comparisons test, *p<0.05). (I) Microarray analysis comparing gene expression after 6h 10μM auxin treatment and auxin+DEX treatment. P=leaf primordia. S=SAM. White arrows and inset image in C and D highlight PIN1-GFP polar localisation in primordia. White arrows in F and G highlight *DR5rev::GFP* expression in the vasculature of developing leaves. Values are means ± SE.

To test whether *ago10^zwl-3^* mutants may have altered auxin transport we investigated the effect of the polar auxin transport inhibitor naphthylphthalamic acid (NPA) on *ago10^zwl^*^-3^ SAM development. NPA increased the frequency of CUP and in particular EA phenotypes. In contrast, NPA treatment decreased the frequency of WT and in particular PIN phenotypes (Fig. 5E). It is also worth noting that NPA had a stronger effect on the frequency of PIN, CUP and EA phenotypes than WT phenotypes. One interpretation of the data is that *ago10^zwl-3^* with mutant phenotypes have impaired auxin transport because they are highly sensitive to further inhibition of polar auxin transport by NPA.

We hypothesised the *ind* mutation might affect *ago10^zwl-3^* sensitivity to NPA because *ago10^zwl-3^* mutants had increased *IND* expression and ectopic expression of IND strongly reduced PIN-GFP levels and polar localisation. Compared to *ago10^zwl-3^* mutants, the *ago10^zwl^*^-3^ *ind-6* mutant was almost completely insensitive to NPA treatment (Fig. 5E). This demonstrates that NPAs effect on *ago10^zwl-3^* was dependent on IND function.

We then tested whether ectopic expression of IND-GR would affect auxin signalling. We observed the level of the auxin-signalling reporter, *DR5rev::GFP*, after DEX treatment of IND-GR lines using confocal microscopy. In the mock-treated control, the DR5rev::GFP signal was detected in the meristem and in the leaf primordia (Fig. 5F). After IND induction, DR5rev::GFP signal was reduced in the meristem suggesting IND represses auxin signalling in this tissue (Fig. 5G).

Since *ago10^zwl-3^* mutants are defective in auxin levels in the embryo [22], we tested whether exogenous auxin treatment can affect the frequency of *ago10^zwl-3^* mutant phenotypes. IAA treatment had no significant effect on the frequency of *ago10^zwl-3^* and double mutant phenotypes (Fig. 5H). Therfore this suggest that *ago10^zwl-3^* and double mutant phenotypes are not due to reduced auxin levels.

We have previously shown that IND’s regulation of gene expression is generally auxin-dependent using microarray analysis (E-MTAB-3812) [28], and here we have reanalysed our microarray data to investigate meristem associated gene regulation and auxin-regulated genesets in detail. To investigate the effect of inducing IND expression on global auxin-regulated responses we analysed how IND effects an auxin-regulated geneset. We treated 7 day old IND-GR seedlings with auxin (AUX), DEX or AUX+DEX and measured gene expression changes using microarray analysis. 1129 genes were differentially expressed (p<0.05, fold change >1.5 or <−1.5) by 6h AUX vs. DEX+AUX treatment (10μM). Linear regression analysis shows that compared to auxin treatment alone the addition of DEX (AUX+DEX) generally antagonised auxins effect on gene expression (Fig. 5I, R^2^ = 0.06078, pearson r = −0.7796. P= <0.0001). This shows that inducing IND expression reduces auxin responses in seedlings.

Gene set enrichment analysis (GSEA) analysis showed that IND induction did not significantly effect enrichment of meristem maintenance and meristem initiation gene sets (P value = >0.23). However, IND induction in the presence of auxin did negatively downregulate meristem maintenance and meristem initiation gene sets (P value = <0.01, Additional file 2: Table S1). This suggests that IND in the presence of auxin negatively regulates genes required for meristem maintenance.

We then investigated whether candidate meristem associated genes were mis-regulated in *ago10^zwl-3^* mutants and whether this misregulation was IND dependent. Of the genes we screened (Additional file 1: Figure S3) most were significantly downregulated in *ago10* in at least one of the mutant phenotype groups, which could be due to the reduction of meristem identity. However, loss of *IND* function did not rescue their expression, suggesting IND did not function through the contol of these meristem regulators. However, it is possible that IND may repress CUC1 during SAM development because we noted that CUC1 was conspiciously downregulated in all *ago10^zwl-3^* mutants, and *CUC1* expression was rescued in the *ago10^zwl-3^ ind-6* double mutant (Fig. 6A, Additional file 1: Figure S3). Consistant with this, we found IND repressed *CUC1* in the presense of CHY (Fig. 6B) and bound the *CUC1* promoter directly in ChIP experiments (Fig. 6C).

**Fig. 6.**
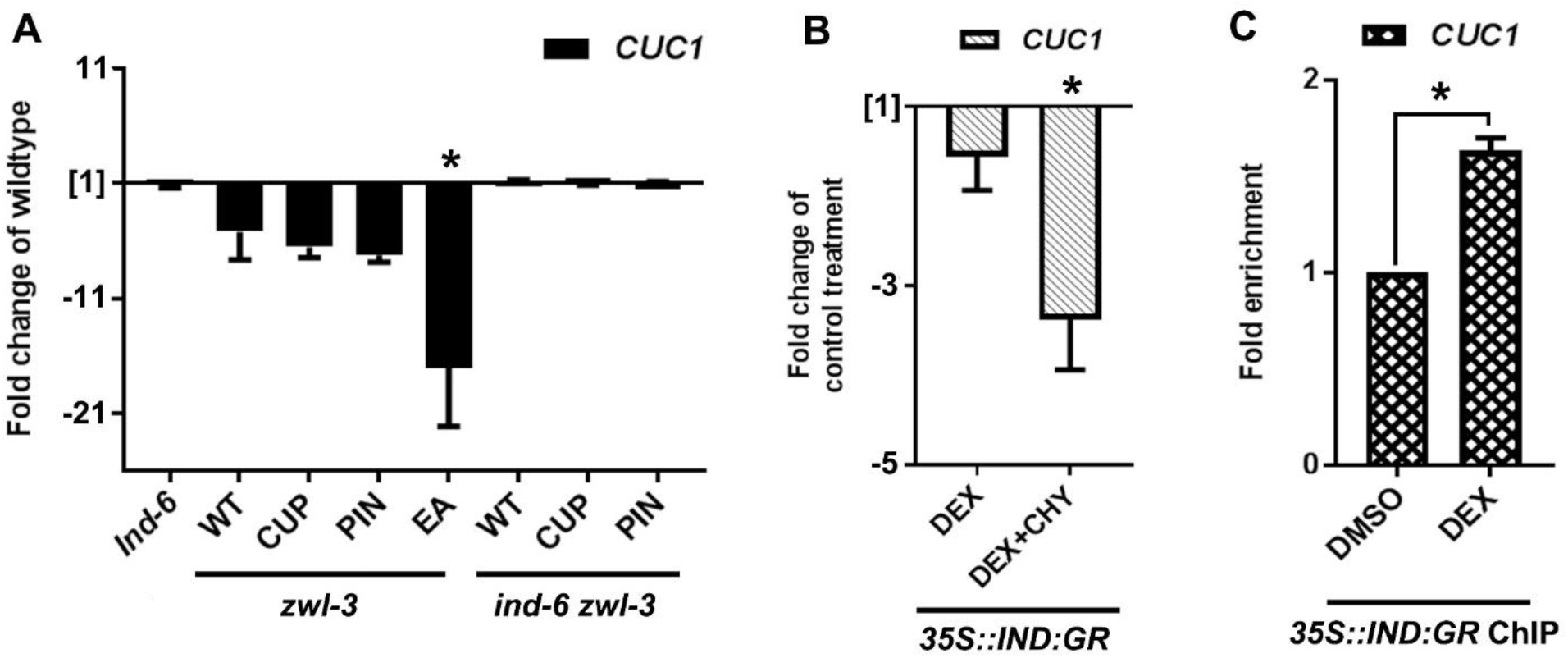
AGO10 regulates *CUC1* expression via direct regulation by *IND*. (A) *CUC1* expression is reduced in *ago10^zwl-3^* mutant and expression is rescued in the *ago10^zwl-3^,ind-6* double mutant measured by qRT-PCR (n=2). (B) qRT–PCR in 7-day-old 35S::IND-GR seedlings after 6h ±DEX and ±cycloheximide (CHY) treatment (n=3). IND directly downregulates *CUC1* expression. (C) ChIP using anti-GR antibody followed by qPCR using *35S::IND:GR* line (n=3). IND-GR binds an upstream element in the *CUC1* promoter (upstream 29 bp-34 bp 5’-UTR) that encodes a putative IND binding site (n=3). *p<0.05 two tailed t test. Values are means ± SE.

## Discussion

During embryo development AGO10 is required to maintain the stem cell niche and prevent meristem termination after germination [8, 10, 11, 14]. The direct functions of AGO10 are well characterised. In particular, AGO10 maintains *HD-ZIP III* expression by sequestering miRNA 165/166 [13]. Several indirect downstream functions have also been identified that are important for signal transduction and feedback regulation [16, 33, 39, 40]. For example, in the embryo *AGO10* represses auxin signalling and this requires *ARF2* [22]. However, a direct signal transduction pathway linking AGO10 and auxin responses has not been established. We found *IND* is a missing link integrating *AGO10* function with auxin signalling during SAM development. We suggest the major function of *AGO10* is to repress *IND* expression to prevent meristem termination and organ fusions in the SAM and regulate fruit development.

### AGO10 and indehiscent function in a network to regulate SAM development

Our work, together with previous findings, supports our proposition that *AGO10* and *IND* function in a network together with *HD-ZIP IIIs* and *SPT* to regulate SAM development (Fig 7). We show several lines of evidence that *IND* functions downstream of *AGO10* to maintain SAM development. Firstly, AGO10 and IND were expressed in similar tissues. Secondly, *IND* expression was increased in *ago10* mutants, both in seedlings and in mature fruit. Thirdly, increasing IND expression phenocopied the *ago10^zwl-3^* mutant SAM phenotype and conversely, reducing *IND* expression in the *ago10^zwl-3^,ind-6* double mutant partially restored the *ago10^zwl-3^* SAM and fruit phenotypes. A surprising observation was that loss of IND function did not rescue the CUP phenotype in *ago10^zwl-3^,ind-6* double mutants even though ectopic expression of IND can induce CUP phenotypes. This suggests the *ago10^zwl-3^* CUP phenotypes are caused by a molecularly distinct pathway that is IND independent. An alternative hypothesis is that the *ago10^zwl-3^* phenotypes are progressively more severe where for example WT<CUP<PIN<EA. Perhaps in the double mutant some CUP phenotypes are rescued to become WT or PIN phenotypes but frequency of CUP phenotypes do not change because the PIN and EA phenotypes become CUP.

**Fig. 7.**
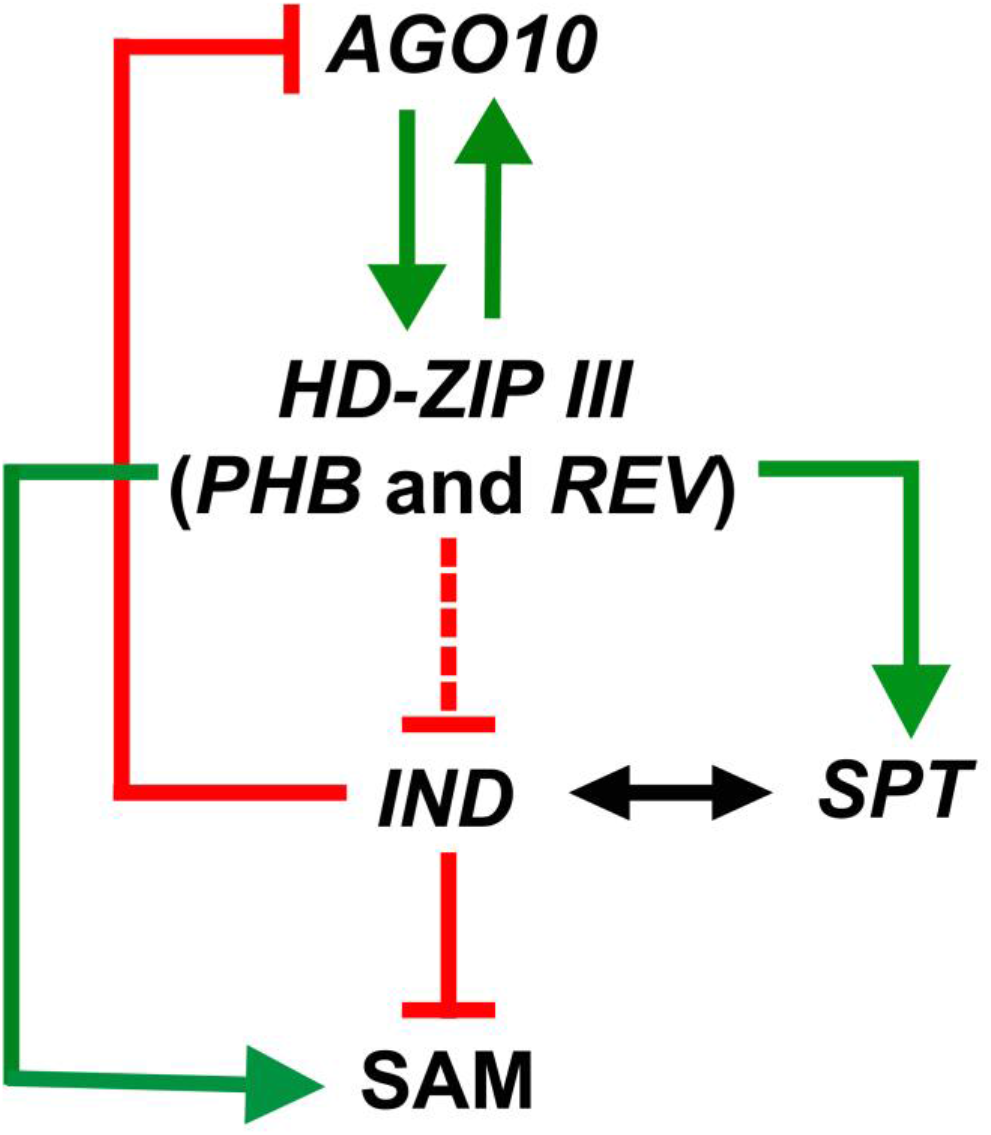
Proposed genetic pathway regulating SAM development. Red lines represent repressive and green arrows are activating regulation [8, 10, 19, 37, 53]. Double arrowed line represent protein interaction [24]. Solid lines represent direct regulation and dotted lines represent indirect regulation.

What is the molecular mechanism regulating *IND* by AGO10? We suggest *IND* is regulated downstream of AGO10 via a signal transduction pathway involving HD-ZIP III’s and SPT transcription factors. *IND* is repressed downstream of HD-ZIP III’s, as we found *IND* expression was induced in *phb*, *phv* and *rev* mutants and overexpression of PHB and REV repressed *IND*. The repression of *IND* by PHB and REV is likely to be indirect, because repression was lost when protein synthesis was blocked. This is consistent with the finding that PHB, PHV and REV were not found to bind IND in genome wide binding studies [37, 38].

We then propose AGO10 may modulate IND activity by inducing *SPT* via direct regulation by HD-ZIP IIIs. To support this conclusion we show PHB and REV directly induces *SPT* as we show PHB and REV can induce SPT even in when protein synthesis was blocked, and REV and PHV was found to bind *SPT* in genome wide binding studies [37, 38]. Consistent with this, we found SPT expression was repressed when AGO10 function is knocked out in mutants. IND heterodimerises with SPT to regulate gene expression and is required for formation of radialised tissues in the SAM and fruit [23, 24]. The reduced expression of *SPT* in *ago10^zwl-3^* mutants may represent a compensatory mechanism to limit IND activity. In WT plants SPT is expressed in the stem cell niche and is required to maintain meristem size [41, 42], therefore regulation by HD-ZIP IIIs may ensure proper tissue localisation of *SPT* which could be important for localising IND function. The weak expression of IND-YFP in the meristem limited our analysis of tissue specific expression in this study, but more sensitive methods for analysing expression such as GFP fusion constructs, FACS, or laser dissection of meristem tissues could provide this data in the future.

IND may also feedback to regulate AGO10 and miR166 expression. However, we did not observe increased expression of *AGO10* in *ind* mutants, which suggests that other factors regulate AGO10 in an *ind* background or the regulation of *AGO10* by ectopic expression of IND-GR is an off-target effect.

Our data suggests AGO10 regulates *IND* to control auxin signalling and expression of genes associated with meristem function. During embryo development AGO10 reduces auxin signalling and this indirectly involves ARF2 [22]. However, a direct signalling pathway from AGO10 to regulating auxin responses pre or post embryogenesis has not been described. We were not able to directly monitor auxin responses in *ago10^zwl-3^* mutants because fluorescent auxin response and transport markers were not available in the L*er* background. However, several lines of evidence suggest AG010 is important for maintaining auxin responses via *IND*. Firstly, auxin transport may be altered in *ago10^zwl-3^* mutants because inhibiting auxin transport strongly increases meristem termination phenotypes in *ago10^zwl-3^*, and this response is completely dependent on *IND* function. Secondly, we also observed that overexpression of IND globally reduced auxin responsive gene expression. Thirdly, since IND directly binds auxin to regulate PID [28], we suggest that AGO10 regulates *IND* levels to maintain proper expression of *PID*. Consistent with this hypothesis, ectopic expression of IND directly reduced the levels and polar localisation of PIN1 and reduced auxin signalling in the meristem.

AGO10 may regulate SAM development by maintaining *CUC1* expression [8, 43, 44]. In the *ago10* mutant IND overexpression directly downregulates *CUC1* expression, which could cause loss of specification of the boundary zone and meristem identity [45]. One prediction, yet to be tested, is that overexpression of *CUC1* would rescue *ago10* mutant phenotypes.

Organ fusion/separation is an important process to regulate development in animals and plants [46]. Our analysis of post-embryonic development of *ago10* mutants reveal its role in regulating organ fusion/separation in the fruit and SAM. AGO10 prevents fusion of the valve margins during fruit development, and also leaf primordia fusion during SAM development by repressing *IND*. During fruit development *IND* with *SPT* and *SPT* with *CUC1* function together to seal the top of the fruit by promoting post-genital fusion of the fruit apex (style tissue) during gynaecium development [47]. Therefore, we suggest a general function of *IND*, together with *CUC1* and *SPT*, is to promote organ fusions.

It is probable that fruit evolved from modified leaves and SAMs [48] as many genes regulating SAM development also regulate fruit development, and our work identifying a conserved mechanism regulating SAM and fruit development further supports this conclusion.

## Conclusion

In this study, we have shown that AGO10 regulates auxin responses via a signal transduction cascade involving *IND* (Fig. 7). This pathway is conserved during both SAM and the fruit development. It will be interesting to investigate whether post-embryonic SAM development is regulated by the interaction of IND, ETTIN and auxin, as it is during fruit development. We provide a genetic framework to test whether the AGO10-IND pathway regulates other processes that have been linked to these genes such as cytokinin signalling, senescence, and organ size.

## Methods

### Plant growth and materials

Seeds were sown on Levington^®^ compost and stratified at 4°C for three days. Plants were illuminated for 16 hours with light delivered at 120μmol m-2 sec-1 at a constant temperature of 23°C in a Versatile Environmental Test Chamber MLR 350-HT (Panasonic, Japan). Distilled water was used for watering seeds in order to control the nutrient supplementation. For growth on agar, seeds were surface-sterilized in 70% ethanol for 10 minutes then treated with 10% bleach, 0.1% (v/v) Triton X-100 for 5 minutes, and finally washed three times with autoclaved water. After stratification at 4°C for three days, the sterile seeds were sown on 0.8% agar supplemented with ½ Murashige and Skoog salts (Murashige and Skoog, 1962) plus vitamins (MS; Duchefa Biochemie, M0222) and 0.5% (w/v) glucose (D-(+)-Glucose, Sigma Aldrich, G7021) in sterile plates. Plates were sealed with micropore tape to maintain sterility while allowing gas exchange. For growth in liquid culture, sterile seeds were sown in 10mL 0.5 % MS medium in a 50mL Falcon tube. Tubes were constantly illuminated in light delivered at 120μmol m-2 sec-1 at a constant temperature of 23°C, and aerated by shaking upright at 60 rotations per minute (rpm). Mutant and transgenic lines *ind-6, zwl-3, ind-6 zwl-3, pZLL::YFP-ZLL zll-1* and *35S::REV:GR* were in Landsberg *erecta* (L*er*) background [8, 29, 33, 36], and *ago10-4, phb-12 er-2, phv-11 er-2, rev-6 er-2, 35S::IND:GR, pIND::GUS, 35S:LhGR>>PHB, pIND::IND:YFP, 35S::IND:GR pPIN1::PIN1:GFP* and *35S::IND:GR DR5rev::GFP* were in Col-0 background [13, 18, 19, 26, 28, 35].

### Hormone and chemical treatments

Seedlings were grown in plant agar medium or liquid culture medium containing hormones and chemicals: Indole-3-acetic acid (IAA), N-1-naphthylphthalamic acid (NPA), Cycloheximide (CHY), Dexamethasone (DEX) and mock solutions. Final concentrations of 10μM IAA, 10μM NPA, 10μM DEX and 10μM CHY were used for treatment. The mock solution contained DMSO (Fisher, BP231) and dH2O (Fisher, W/0100/21). All treated plants with their respective controls were grown simultaneously under the same conditions.

### Confocal and standard light microscopy

Analysis of SAM phenotypes was analysed at 3 and 14 days post germination. Seedlings were transferred to a Petri dish filled with sterile water. Forceps were used to hold one cotyledon while pulling the second cotyledon downwards to peel the seedling into two. This peeled cotyledon was transferred to a microscope slide and aligned on top of 1 % agarose gel. Two cotyledons of a seedling were observed under a light microscope to analyse the phenotype of shoot apical meristem. For confocal microscopy a stereomicroscope was used to dissect and analyse the plant material. SAMs were analysed by staining with 5 μg/mL of propidium iodide (PI) solution for 6 hours. The stained samples were mounted on microscope slides and imaged on a confocal microscope. Propidium iodide can be excited by a 514 nm argon laser beam and emits between 580-610 nm. Transgenic embryos or seedlings or fruits (*pAGO10::AGO10:YFP, pPIN1::PIN1:GFP, DR5rev::GFP* and *pIND::IND:YFP*) were mounted on microscope slides with a slab of 1% plant agar and imaged using an Olympus FV1000 confocal microscope. Laser setting was selected and changed using software FV10-ASW. YFP can be excited by a 514 nm argon laser beam and emits between 520-530 nm. GFP can be excited by a 488 nm argon laser beam and emits between 495-515 nm. Chlorophyll excitation was at 488 nm and emission was between 650-710 nm. Captured images were processed using FV10-ASW viewer or Image J.

### Scanning electron microscopy (SEM)

Samples were fixed in 3% Glutaraldehyde/0.1M sodium cacodylate buffer, washed in 0.1M sodium cacodylate buffer to remove unbound fixative and secondarily fixed in 2% aqueous osmium tetroxide for 1 hour. Specimens were dehydrated through a sequentially graded series of ethanol, 50%-100%, for 30 minutes per step, finally into 100% ethanol before being dried over anhydrous copper sulphate. Specimens were critically point dried using CO2 as the transitional fluid. After drying, the specimens were mounted on 12.5mm diameter stubs, attached with sticky tabs and coated in an Edwards S150B sputter coater with approximately 25 - 30 nm of gold. Specimens were viewed using a Philips SEM XL-20 Scanning Electron Microscope at an accelerating voltage of 20kV in Biomedical Science Electron Microscopy Unit, University of Sheffield.

### β-Glucuronidase (GUS) Histology

GUS assay was performed on *pIND::GUS* seedlings at different developmental stages. Samples were vacuum infiltrated and incubated in the GUS assay buffer (0.1M phosphate buffer [pH 7], 10mM EDTA, 0.1% Triton X-100, 1mg/mL X-Glue A, 2mM potassium ferricyanide) overnight at 37°C, and cleared in 50% ethanol. For histology, Samples were rinsed in 70% v/v ethanol and fixed in 100% EtOH: glacial acetic acid (7:1 v/v) at room temperature (19–22°C) overnight until the complete removal of chlorophyll. Samples were embedded for sectioning using Technovit 7100 resin solution (TAAB, #T218) following the manufacturer’s instructions. Sections (8 μm) were taken using a Leica RM2145 microtome. GUS staining was observed under a light microscope and photographs were taken with a CCD camera.

### Quantitative reverse transcriptase PCR (qRT-PCR)

Total nucleic acid (TNA) was extracted using a phenol-chloroform extraction procedure adapted from [49]. Complementary DNA (cDNA) from 1–2 μg of DNase I treated TNA was synthesised using a High Capacity cDNA Reverse Transcription Kit using random primers (Invitrogen, #4374966). qRT-PCR was performed with SYBR Green Jump-start Taq Ready-mix (Sigma, S4438) on the Mx3005P qPCR System (Agilent Technologies Genomics). Reactions were prepared using 2X JumpStart Taq Ready Mix, 1X ROX Reference Dye, 300nM forward primer, 300nM reverse primer, 500ng template DNA and nuclease-free water and 15μl of each reaction was transferred to an optical 96 well plate. The plate was covered with an optical adhesive film (Bio-Rad, #MSB-1001). PCR products were analysed by agarose gel electrophoresis and the disassociation curve analysis to confirm that the PCR primers produced a single product of the correct predicted size. The threshold cycle (CT) was automatically determined by the Mx3005P qPCR System, and comparative CT method (also known as the 2 ^−ΔΔCT^ method) was used to analyse the qRT-PCR data [50]. *ACTIN2* was used as a normalisation control. Primers used for qRT-PCR were listed in Additional file 2: Table S2.

### Chromatin immunoprecipitation (ChIP)

*35S::IND:GR* seeds were grown for 7 days in 50 ml of liquid culture medium with constant shaking. After 7 days of growth under constant light, seedlings were treated with a final concentration of 10 μM DEX (treatment) and DMSO (control) for 6 hours. The ChIP experiments were performed as previously described [26]. Q-PCR was performed with SYBR Green Jump-start Taq Ready-mix (Sigma, S4438) on the Mx3005P qPCR System (Agilent Technologies Genomics) and using the primers Pro CUC1 F, Pro CUC1 R, Pro AGO10 F, and Pro AGO10 R (Additional file 2: Table S2). The values correspond to the fold enrichment between DEX treated input with the GR antibody and DMSO treated input with the GR antibody.

### Geneset enrichment analysis (GSEA)

GSEA is a powerful analytical tool used to study groups of genes or proteins that share common biological function, protein domain, chromosomal location, or regulation in large datasets, GSEA is by the Broad Institute [51]. *Arabidopsis thaliana* GO library files was prepared using gene ontology consortium *Arabidopsis thaliana* GO annotations [52]. One thousand sample permutations were selected for any analysis. Normalized Enrichment Score (NES) was used to compare analysis results across gene sets. GSEA report was viewed in a web browser (HTML Report) and transferred to Excel.

## Declarations

### Ethics approval and consent to participate

Not applicable

## Consent for publication

Not applicable

## Availability of data and material

The datasets generated analysed during the current study are available in the ArrayExpress repository (E-MTAB-3812). Other public datasets GSE26722 and GSM1925338 were also analysed.

## Author contributions

MV and KS performed all experiments. KS supervised the project. KS and MV wrote the manuscript. All authors read and approved the final manuscript.

## Acknowledgements

We thank Prof Thomas Laux for kindly providing the *pZLL::YFP-ZLL zll-1* line, Prof Miltos Tsiantis for kindly providing the *35S:LhGR>>PHB* line and Prof Lars Ostergaard for kindly providing the *pIND::IND:YFP* line. We thank Dr Chris Hill for processing samples for SEM at the University of Sheffield, Giulia Arsuffi for technical assistance, Matt parker for assistance with GSEA Arabidopsis thaliana GO library file preparation and Dr Irina-ioana Mohorianu for processing the RNAseq data. We also thank Prof Andrew Flemming for critical analysis of the manuscript.

## Open Access

This article is distributed under the terms of the Creative Commons Attribution 4.0 International License (http://creativecommons.org/licenses/by/4.0/), which permits unrestricted use, distribution, and reproduction in any medium, provided you give appropriate credit to the original author(s) and the source, provide a link to the Creative Commons license, and indicate if changes were made. The Creative Commons Public Domain Dedication waiver (http://creativecommons.org/publicdomain/zero/1.0/) applies to the data made available in this article, unless otherwise stated. All work complied with with local, national and international guidelines and legislation for plant research.

## Abbreviations

SAM: Shoot apical meristem
AGO10^zwl-3^: ZWILLE-3 in Ler background.
AGO10-4: AGO10-4 in Col background.
IND: INDEHISCENT
SPT: SPATULA
YFP: yellow fluorescent protein
GFP: green fluorescent protein
L*er*: Landsberg *erecta*
qRT-PCR: quantitative reverse transcriptase polymerase chain reaction
HD-ZIP III: HOMEODOMAIN LEUCINE ZIPPER class III
REV: REVOLUTA
PHB: PHABULOSA
PHV: PHAVALUTA
PID: PINOID
GUS: beta-Glucuronidase
DEX: dexamethasone
GR: glucocorticoid receptor
ChIP: Chromatin immunoprecipitation
NPA: Naphthylphthalamic acid
GSEA: gene set enrichment analysis.

---

**Fig. S1.**
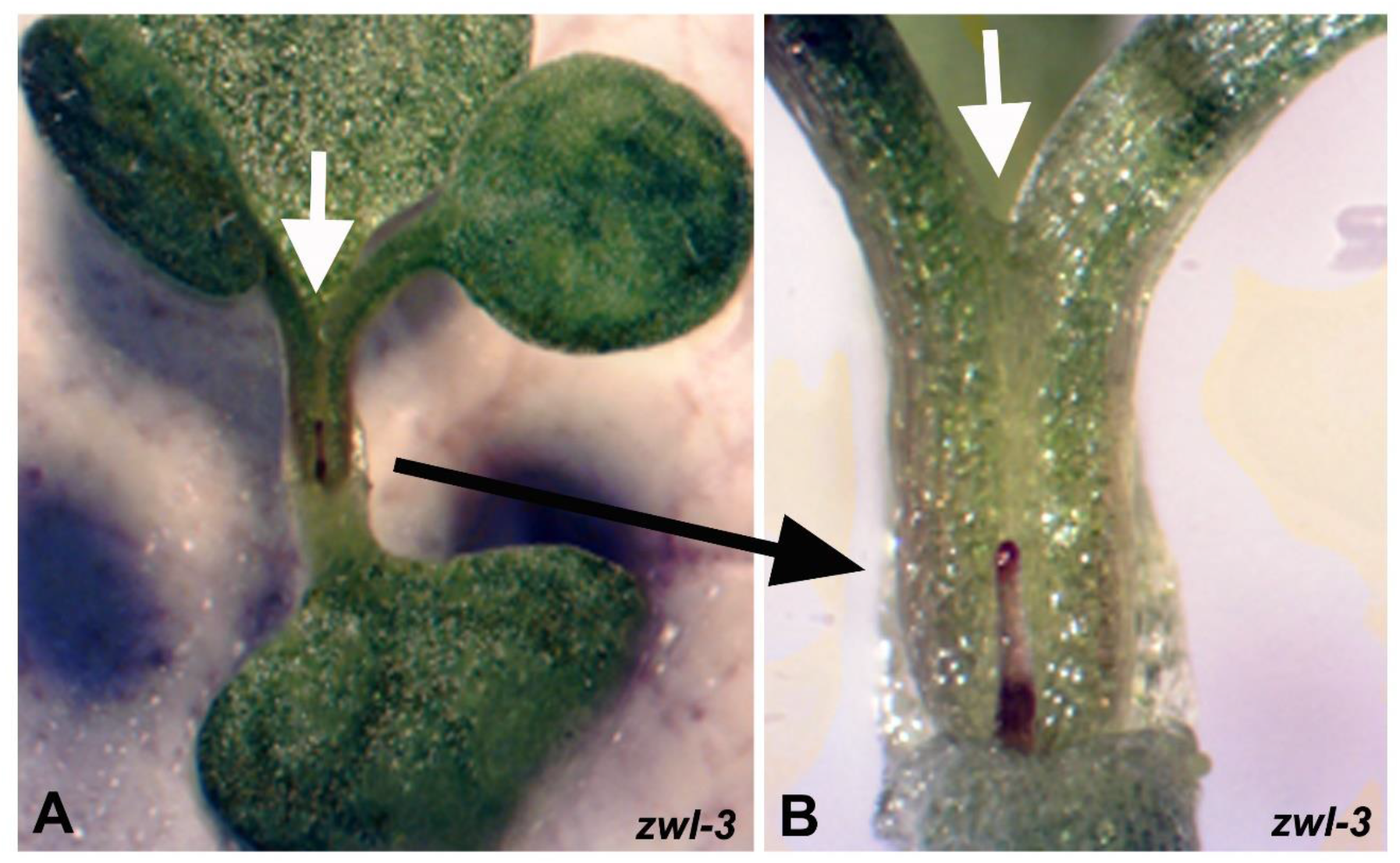
(A) Occasionally (2/91) the first two true leaves are fused in *ago10^zwl-3^* mutants. (B) Higher magnification of fused leaf phenotype. The petioles of the first true leaves are fused together and the meristem appears to have terminated (white arrow). A radialised PIN shaped leaf is observed in place of the 3^rd^ or 4^th^ leaf (black arrow).

**Fig. S2.**
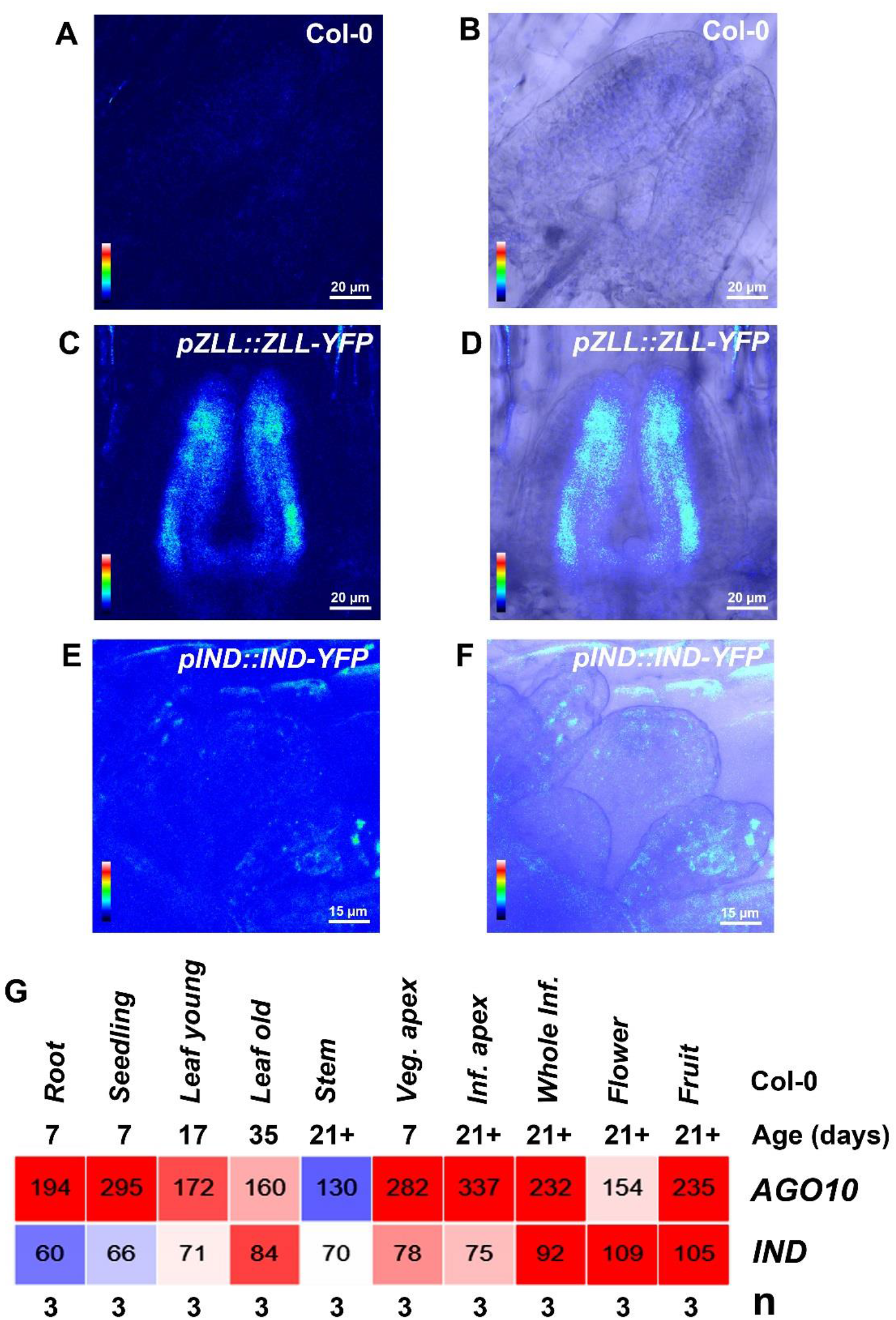
IND is expressed in the meristem and post-embryonic tissues. (A,C,E) False colouring of confocal microscopy image to highlight YFP signal (Rainbow gradient from blue to red represents no expression to high expression). (B,D,F) Merged confocal and light transmission image. (A,B) YFP signal in negative control plant. (C,D) YFP-AGO10 is observed in leaf primordia (E,F) Compared to the negative control, IND-YFP signal is observed in leaf primordia and weakly in the SAM. (G) Absolute signal values of *AG010* and *IND* in different tissues as measured by Tiling expression array dataset [1]. IND signal is highest in the fruit and flower tissues and also in the vegetative meristem (veg. apex).

**Fig. S3.**
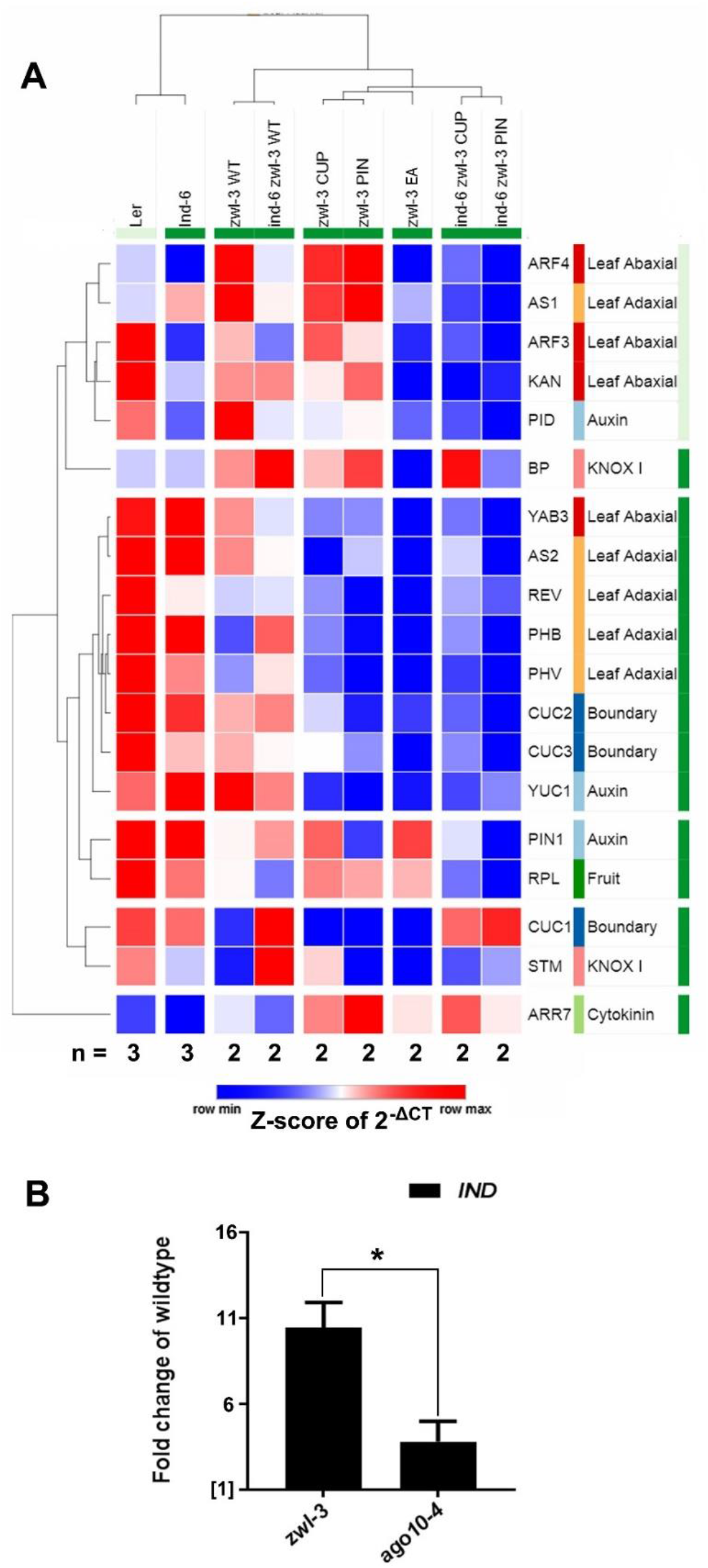
(A) Expression profile of genes associated with meristem development in wild type, *ago10^zwl-3^, ind-6* and double mutants measured by qRT-PCR. (B) *IND* expression is significantly increased in *ago10^zwl-3^* mutants (L*er* background) and to a lesser extent in *ago10-4* mutants (Col background) compared to their wild type backgrounds. *IND* expression is significantly less induced in *ago10-4* mutants compared to *ago10^zwl-3^* mutants (n=3, *ago10^zwl-3^* vs. *ago10-4*, two tailed t-test *p<0.05).

**Fig. S4.**
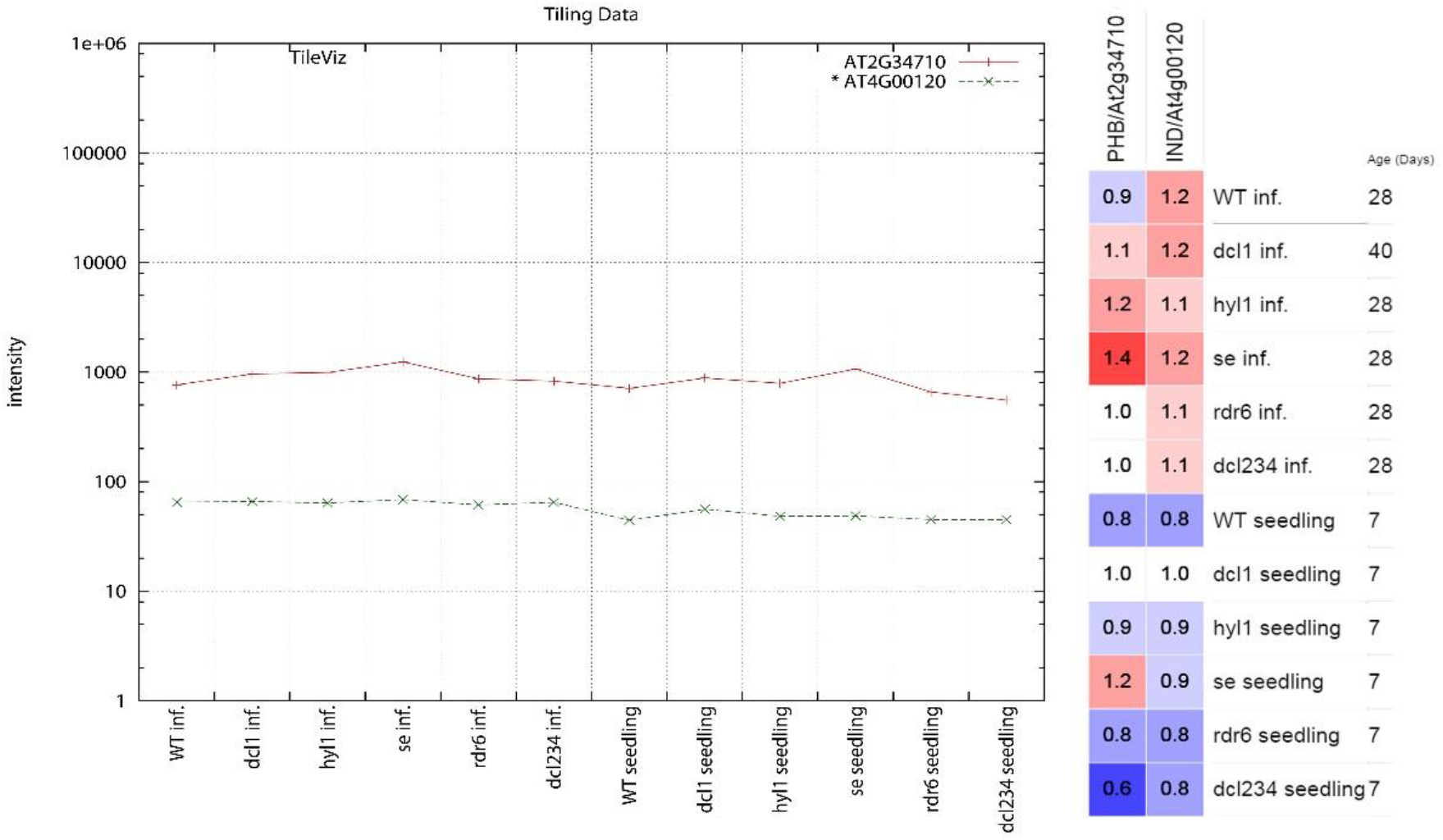
*PHB* and *IND* gene expression in mutants impaired in small RNA biogenesis [2]. On the left line graph shows Intensity values of *PHB* or *IND* in influorescene and seedlings. On the right the heat map displays mean-normalised values of PHB or IND. When compared to wild type, *PHB* expression increased in *se* mutants as shown by Grigg *et al*, [3]. *IND* expression did not change dramatically in the inflorescence or seedling samples of wild type and mutants.

**Fig. S5.**
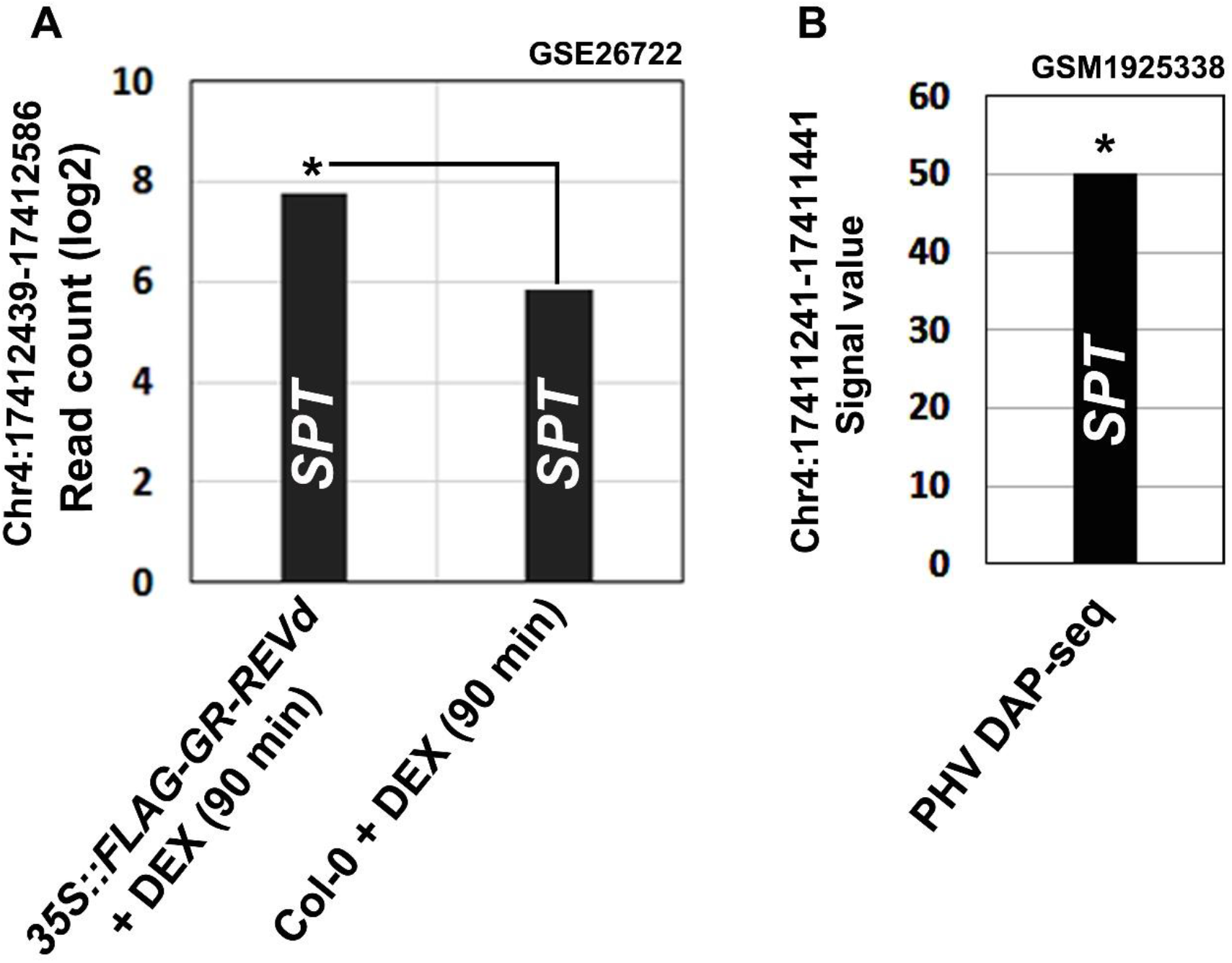
PHV and REV directly bind to *SPT* promoter. PHV DAP-seq (GEO:GSM1925338) [4] and REV ChIP-seq (GEO:GSE26722) [5] data were analysed to identify PHV and REV binding *SPT* promoter. (A) REV can directly bind to *SPT* promoter (Chr4:17412439-17412586). (B) PHV can directly bind to *SPT* promoter (Chr4:17411241-17411441).

**Fig. S6.**
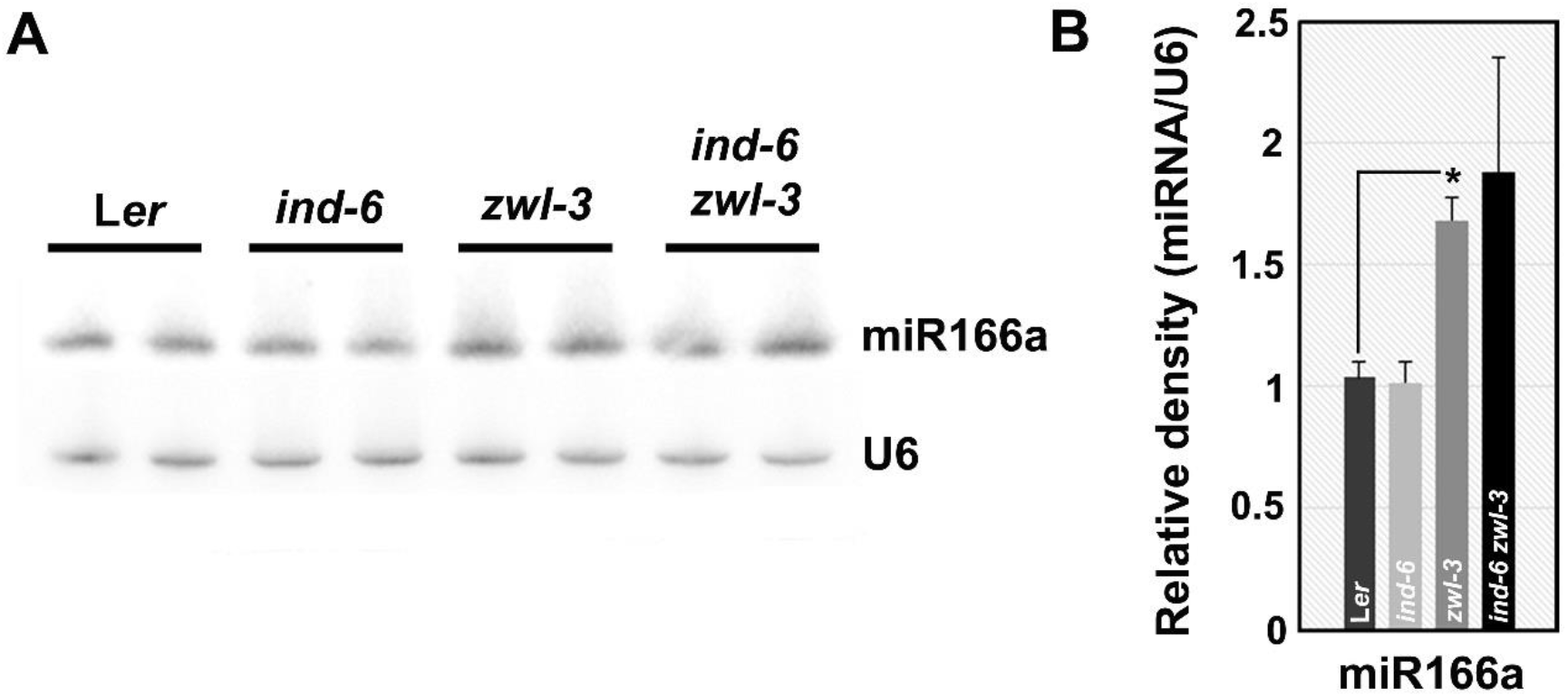
(A) The northern blot image show increased miR166a expression in *ago10^zwl 3^, ind-6 and ind-6 ago10^zwl-3^* double mutants. *U6* is shown as loading control (n=2 biological replicates). (B) Quantification of band intensity form northern blot normalised to *U6* loading control (n=2, two tailed t-test *p<0.05).

**Fig. S7.**
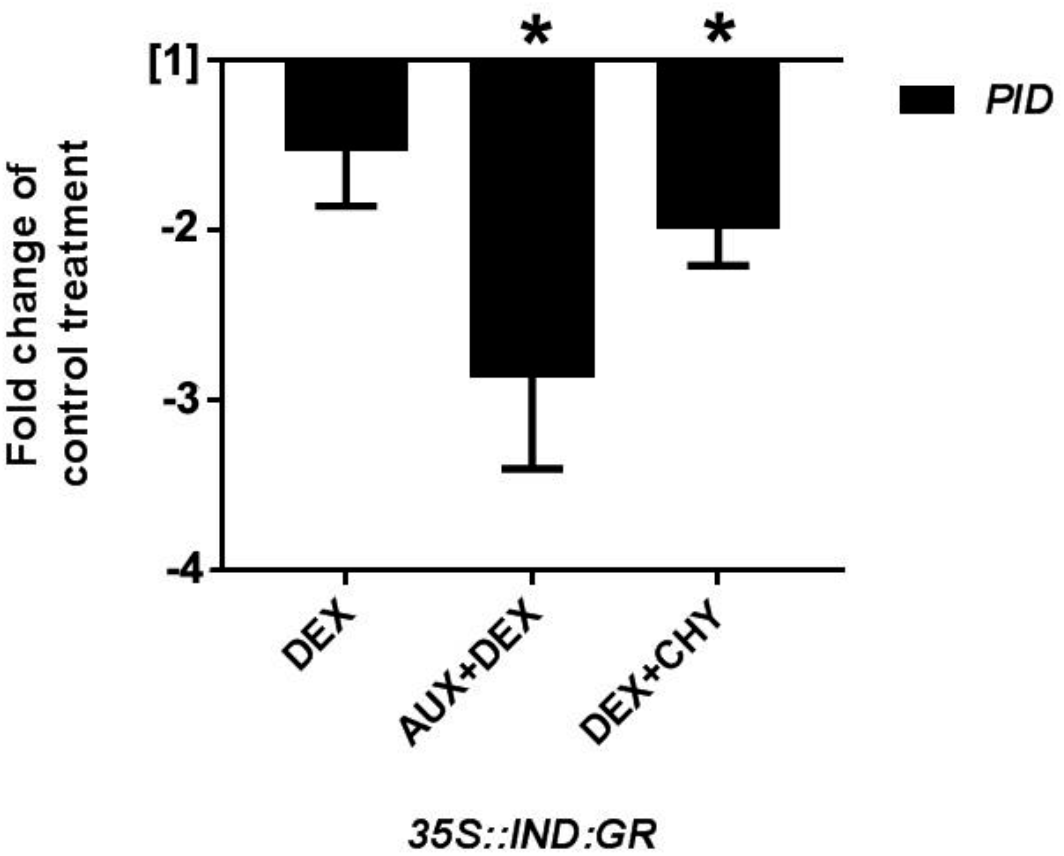
IND directly regulates *PID* expression. *35S::IND:GR* seedlings were treated with DEX, Auxin + DEX (AUX+DEX) or DEX + Cycloheximide (DEX+CHY) for 6h followed by qRT-PCR using *PID* specific primers. Concentrations used were DEX 10μM, Auxin 10μM and CHY 10μM. (n=3, DEX vs DMSO, AUX+DEX vs AUX, DEX+CHY vs CHY, two tailed t-test *p<0.05).

**Table S1:** From GSEA analysis, meristem maintenance and meristem initiation gene sets were significantly negatively enriched in *35S::IND:GR* (AUX+DEX vs. AUX) (E-MTAB-3812).

| GO Biological Process Geneset | Size | DEX vs. DMSO |  | AUX+DEX vs. AUX |  |
| --- | --- | --- | --- | --- | --- |
|  |  | NES | NOM P-val | NES | NOM P-val |
| **Meristem Maintenance** | 67 | -1.13 | 0.235 | -2.09 | <0.0001 |
| **Meristem Initiation** | 123 | -0.64 | 1.000 | -1.40 | 0.007 |

**Table S2:**
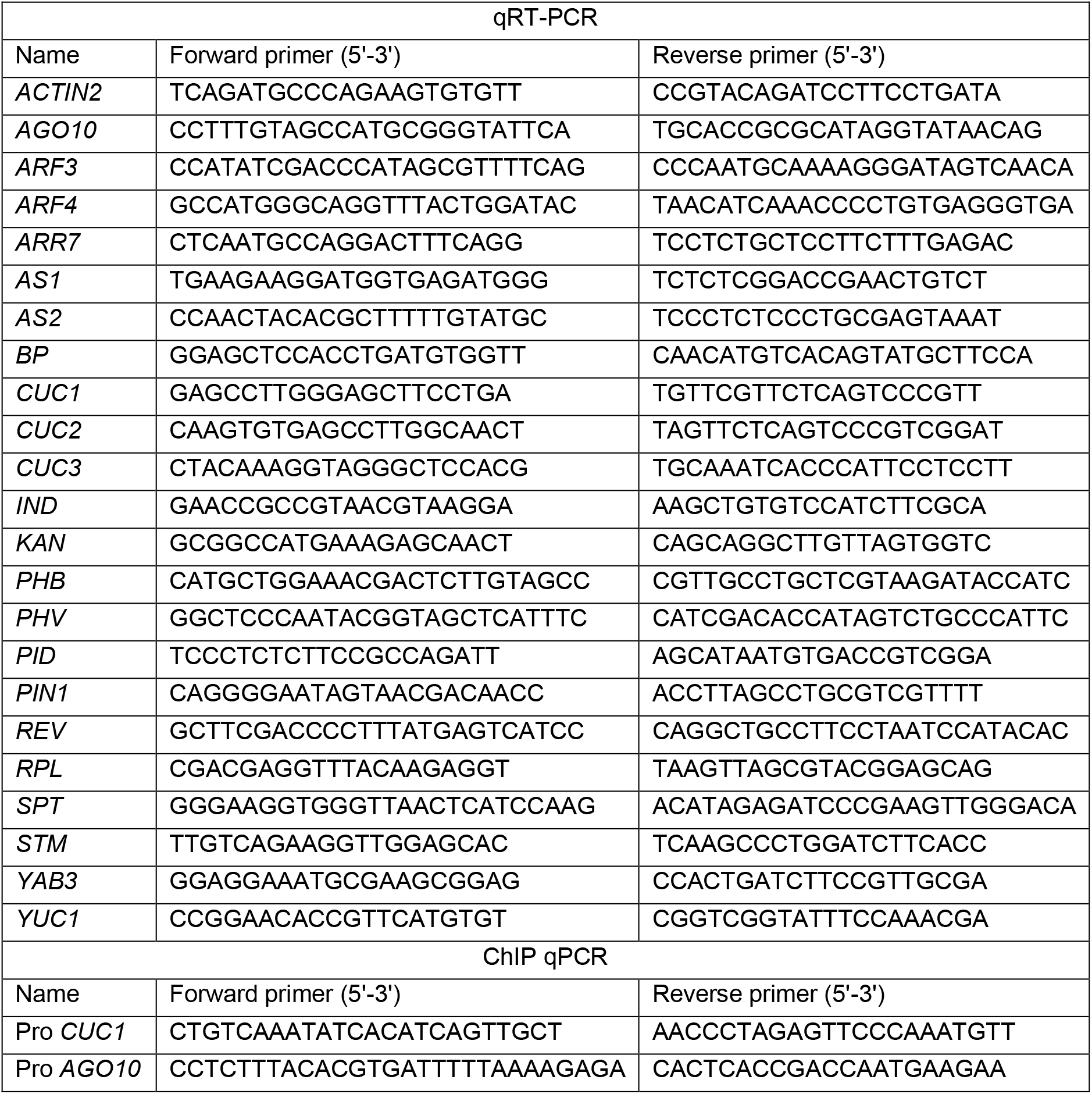
List of primers used for ChIP qPCR and qRT-PCR.

## References

1. Gaillochet C, Daum G, Lohmann JU: O cell, where art thou? The mechanisms of shoot meristem patterning. Current opinion in plant biology 2015, 23:91–97.

2. ten Hove CA, Lu KJ, Weijers D: Building a plant: cell fate specification in the early Arabidopsis embryo. Development 2015, 142(3):420–430.

3. Barton MK: Twenty years on: the inner workings of the shoot apical meristem, a developmental dynamo. Developmental biology 2010, 341(1):95–113.

4. Schoof H, Lenhard M, Haecker A, Mayer KFX, Jurgens G, Laux T: The stem cell population of Arabidopsis shoot meristems is maintained by a regulatory loop between the CLAVATA and WUSCHEL genes. Cell 2000, 100(6):635–644.

5. Aida M, Ishida T, Fukaki H, Fujisawa H, Tasaka M: Genes involved in organ separation in Arabidopsis: an analysis of the cup-shaped cotyledon mutant. The Plant cell 1997, 9(6):841–857.

6. Takada S, Hibara K, Ishida T, Tasaka M: The CUP-SHAPED COTYLEDON1 gene of Arabidopsis regulates shoot apical meristem formation. Development 2001, 128(7):1127–1135.

7. Hibara K, Karim MR, Takada S, Taoka KI, Furutani M, Aida M, Tasaka M: Arabidopsis CUP-SHAPED COTYLEDON3 regulates postembryonic shoot meristem and organ boundary formation. Plant Cell 2006, 18(11):2946–2957.

8. Endrizzi K, Moussian B, Haecker A, Levin JZ, Laux T: The SHOOT MERISTEMLESS gene is required for maintenance of undifferentiated cells in Arabidopsis shoot and floral meristems and acts at a different regulatory level than the meristem genes WUSCHEL and ZWILLE. Plant Journal 1996, 10(6):967–979.

9. Lynn K, Fernandez A, Aida M, Sedbrook J, Tasaka M, Masson P, Barton MK: The PINHEAD/ZWILLE gene acts pleiotropically in Arabidopsis development and has overlapping functions with the ARGONAUTE1 gene. Development 1999, 126(3):469–481.

10. Mcconnell JR, Barton MK: Effect of Mutations in the Pinhead Gene of Arabidopsis on the Formation of Shoot Apical Meristems. Dev Genet 1995, 16(4):358–366.

11. Moussian B, Schoof H, Haecker A, Jurgens G, Laux T: Role of the ZWILLE gene in the regulation of central shoot meristem cell fate during Arabidopsis embryogenesis. The EMBO journal 1998, 17(6):1799–1809.

12. Ji L, Liu X, Yan J, Wang W, Yumul RE, Kim YJ, Dinh TT, Liu J, Cui X, Zheng B et al: ARGONAUTE10 and ARGONAUTE1 regulate the termination of floral stem cells through two microRNAs in Arabidopsis. PLoS genetics 2011, 7(3):e1001358.

13. Zhu H, Hu F, Wang R, Zhou X, Sze SH, Liou LW, Barefoot A, Dickman M, Zhang X: Arabidopsis Argonaute10 specifically sequesters miR166/165 to regulate shoot apical meristem development. Cell 2011, 145(2):242–256.

14. Zhou Y, Honda M, Zhu H, Zhang Z, Guo X, Li T, Li Z, Peng X, Nakajima K, Duan L et al: Spatiotemporal sequestration of miR165/166 by Arabidopsis Argonaute10 promotes shoot apical meristem maintenance. Cell Rep 2015, 10(11):1819–1827.

15. Zhou GK, Kubo M, Zhong R, Demura T, Ye ZH: Overexpression of miR165 affects apical meristem formation, organ polarity establishment and vascular development in Arabidopsis. Plant & cell physiology 2007, 48(3):391–404.

16. Yu Y, Ji LJ, Le BH, Zhai JX, Chen JY, Luscher E, Gao L, Liu CY, Cao XF, Mo BX et al: ARGONAUTE10 promotes the degradation of miR165/6 through the SDN1 and SDN2 exonucleases in Arabidopsis. Plos Biol 2017, 15(2).

17. McConnell JR, Emery J, Eshed Y, Bao N, Bowman J, Barton MK: Role of PHABULOSA and PHAVOLUTA in determining radial patterning in shoots. Nature 2001, 411(6838):709–713.

18. Otsuga D, DeGuzman B, Prigge MJ, Drews GN, Clark SE: REVOLUTA regulates meristem initiation at lateral positions. Plant Journal 2001, 25(2):223–236.

19. Prigge MJ, Otsuga D, Alonso JM, Ecker JR, Drews GN, Clark SE: Class III homeodomain-leucine zipper gene family members have overlapping, antagonistic, and distinct roles in Arabidopsis development. Plant Cell 2005, 17(1):61–76.

20. Emery JF, Floyd SK, Alvarez J, Eshed Y, Hawker NP, Izhaki A, Baum SF, Bowman JL: Radial patterning of Arabidopsis shoots by class III HD-ZIP and KANADI genes. Current biology: CB 2003, 13(20):1768–1774.

21. Mallory AC, Reinhart BJ, Jones-Rhoades MW, Tang G, Zamore PD, Barton MK, Bartel DP: MicroRNA control of PHABULOSA in leaf development: importance of pairing to the microRNA 5’ region. The EMBO journal 2004, 23(16):3356–3364.

22. Roodbarkelari F, Du F, Truernit E, Laux T: ZLL/AGO10 maintains shoot meristem stem cells during Arabidopsis embryogenesis by down-regulating ARF2-mediated auxin response. BMC Biol 2015, 13:74.

23. Moubayidin L, Ostergaard L: Dynamic control of auxin distribution imposes a bilateral-to-radial symmetry switch during gynoecium development. Current biology: CB 2014, 24(22):2743–2748.

24. Girin T, Paicu T, Stephenson P, Fuentes S, Korner E, O’Brien M, Sorefan K, Wood TA, Balanza V, Ferrandiz C et al: INDEHISCENT and SPATULA interact to specify carpel and valve margin tissue and thus promote seed dispersal in Arabidopsis. Plant Cell 2011, 23(10):3641–3653.

25. Groszmann M, Bylstra Y, Lampugnani ER, Smyth DR: Regulation of tissue-specific expression of SPATULA, a bHLH gene involved in carpel development, seedling germination, and lateral organ growth in Arabidopsis. Journal of experimental botany 2010, 61(5):1495–1508.

26. Sorefan K, Girin T, Liljegren SJ, Ljung K, Robles P, Galvan-Ampudia CS, Offringa R, Friml J, Yanofsky MF, Ostergaard L: A regulated auxin minimum is required for seed dispersal in Arabidopsis. Nature 2009, 459(7246):583–U114.

27. Liljegren SJ, Roeder AH, Kempin SA, Gremski K, Ostergaard L, Guimil S, Reyes DK, Yanofsky MF: Control of fruit patterning in Arabidopsis by INDEHISCENT. Cell 2004, 116(6):843–853.

28. Simonini S, Deb J, Moubayidin L, Stephenson P, Valluru M, Freire-Rios A, Sorefan K, Weijers D, Friml J, Østergaard L: A noncanonical auxin-sensing mechanism is required for organ morphogenesis in Arabidopsis. Genes & development 2016, 30(20):2286–2296.

29. Wu H, Mori A, Jiang X, Wang Y, Yang M: The INDEHISCENT protein regulates unequal cell divisions in Arabidopsis fruit. Planta 2006, 224(4):971–979.

30. Arnaud N, Girin T, Sorefan K, Fuentes S, Wood TA, Lawrenson T, Sablowski R, Ostergaard L: Gibberellins control fruit patterning in Arabidopsis thaliana. Genes & development 2010, 24(19):2127–2132.

31. Girin T, Stephenson P, Goldsack CM, Kempin SA, Perez A, Pires N, Sparrow PA, Wood TA, Yanofsky MF, Ostergaard L: Brassicaceae INDEHISCENT genes specify valve margin cell fate and repress replum formation. The Plant journal : for cell and molecular biology 2010, 63(2):329–338.

32. Roeder AH, Ferrandiz C, Yanofsky MF: The role of the REPLUMLESS homeodomain protein in patterning the Arabidopsis fruit. Current biology : CB 2003, 13(18):1630–1635.

33. Tucker MR, Hinze A, Tucker EJ, Takada S, Jurgens G, Laux T: Vascular signalling mediated by ZWILLE potentiates WUSCHEL function during shoot meristem stem cell development in the Arabidopsis embryo. Development 2008, 135(17):2839–2843.

34. Laubinger S, Zeller G, Henz SR, Sachsenberg T, Widmer CK, Naouar N, Vuylsteke M, Scholkopf B, Ratsch G, Weigel D: At-TAX: a whole genome tiling array resource for developmental expression analysis and transcript identification in Arabidopsis thaliana. Genome biology 2008, 9(7):R112.

35. Dello Ioio R, Galinha C, Fletcher AG, Grigg SP, Molnar A, Willemsen V, Scheres B, Sabatini S, Baulcombe D, Maini PK et al: A PHABULOSA/cytokinin feedback loop controls root growth in Arabidopsis. Current biology: CB 2012, 22(18):1699–1704.

36. Wenkel S, Emery J, Hou BH, Evans MM, Barton MK: A feedback regulatory module formed by LITTLE ZIPPER and HD-ZIPIII genes. Plant Cell 2007, 19(11):3379–3390.

37. Brandt R, Salla-Martret M, Bou-Torrent J, Musielak T, Stahl M, Lanz C, Ott F, Schmid M, Greb T, Schwarz M et al: Genome-wide binding-site analysis of REVOLUTA reveals a link between leaf patterning and light-mediated growth responses. The Plant journal : for cell and molecular biology 2012, 72(1):31–42.

38. O’Malley RC, Huang SS, Song L, Lewsey MG, Bartlett A, Nery JR, Galli M, Gallavotti A, Ecker JR: Cistrome and Epicistrome Features Shape the Regulatory DNA Landscape. Cell 2016, 165(5):1280–1292.

39. Tucker MR, Roodbarkelari F, Truernit E, Adamski NM, Hinze A, Lohmuller B, Wurschum T, Laux T: Accession-specific modifiers act with ZWILLE/ARGONAUTE10 to maintain shoot meristem stem cells during embryogenesis in Arabidopsis. BMC Genomics 2013, 14:809.

40. Liu Q, Yao X, Pi L, Wang H, Cui X, Huang H: The ARGONAUTE10 gene modulates shoot apical meristem maintenance and establishment of leaf polarity by repressing miR165/166 in Arabidopsis. The Plant journal : for cell and molecular biology 2009, 58(1):27–40.

41. Schuster C, Gaillochet C, Medzihradszky A, Busch W, Daum G, Krebs M, Kehle A, Lohmann JU: A regulatory framework for shoot stem cell control integrating metabolic, transcriptional, and phytohormone signals. Developmental cell 2014, 28(4):438–449.

42. Gaillochet C, Stiehl T, Wenzl C, Ripoll JJ, Bailey-Steinitz LJ, Li L, Pfeiffer A, Miotk A, Hakenjos JP, Forner J et al: Control of plant cell fate transitions by transcriptional and hormonal signals. Elife 2017, 6.

43. Spinelli SV, Martin AP, Viola IL, Gonzalez DH, Palatnik JF: A mechanistic link between STM and CUC1 during Arabidopsis development. Plant physiology 2011, 156(4):1894–1904.

44. Aida M, Ishida T, Tasaka M: Shoot apical meristem and cotyledon formation during Arabidopsis embryogenesis: interaction among the CUP-SHAPED COTYLEDON and SHOOT MERISTEMLESS genes. Development 1999, 126(8):1563–1570.

45. Furutani M, Vernoux T, Traas J, Kato T, Tasaka M, Aida M: PIN-FORMED1 and PINOID regulate boundary formation and cotyledon development in Arabidopsis embryogenesis. Development 2004, 131(20):5021–5030.

46. Ray HJ, Niswander L: Mechanisms of tissue fusion during development. Development 2012, 139(10):1701–1711.

47. Nahar MA, Ishida T, Smyth DR, Tasaka M, Aida M: Interactions of CUP-SHAPED COTYLEDON and SPATULA genes control carpel margin development in Arabidopsis thaliana. Plant & cell physiology 2012, 53(6):1134–1143.

48. Girin T, Sorefan K, Ostergaard L: Meristematic sculpting in fruit development. Journal of experimental botany 2009, 60(5):1493–1502.

49. White JL, Kaper JM: A Simple Method for Detection of Viral Satellite Rnas in Small Plant-Tissue Samples. Journal of virological methods 1989, 23(2):83–93.

50. Schmittgen TD, Livak KJ: Analyzing real-time PCR data by the comparative C(T) method. Nature protocols 2008, 3(6):1101–1108.

51. Subramanian A, Tamayo P, Mootha VK, Mukherjee S, Ebert BL, Gillette MA, Paulovich A, Pomeroy SL, Golub TR, Lander ES et al: Gene set enrichment analysis: a knowledge-based approach for interpreting genome-wide expression profiles. Proceedings of the National Academy of Sciences of the United States of America 2005, 102(43):15545–15550.

52. Yi X, Du Z, Su Z: PlantGSEA: a gene set enrichment analysis toolkit for plant community. Nucleic Acids Res 2013, 41 (Web Server issue):W98–103.

53. Brandt R, Xie Y, Musielak T, Graeff M, Stierhof YD, Huang H, Liu CM, Wenkel S: Control of stem cell homeostasis via interlocking microRNA and microProtein feedback loops. Mechanisms of development 2013, 130(1):25–33.

## References

1. Laubinger S, Zeller G, Henz SR, Sachsenberg T, Widmer CK, Naouar N, Vuylsteke M, Scholkopf B, Ratsch G, Weigel D: At-TAX: a whole genome tiling array resource for developmental expression analysis and transcript identification in Arabidopsis thaliana. Genome biology 2008, 9(7):R112.

2. Laubinger S, Zeller G, Henz SR, Buechel S, Sachsenberg T, Wang JW, Ratsch G, Weigel D: Global effects of the small RNA biogenesis machinery on the Arabidopsis thaliana transcriptome. Proceedings of the National Academy of Sciences of the United States of America 2010, 107(41):17466–17473.

3. Grigg SP, Canales C, Hay A, Tsiantis M: SERRATE coordinates shoot meristem function and leaf axial patterning in Arabidopsis. Nature 2005, 437(7061):1022–1026.

4. O’Malley RC, Huang SS, Song L, Lewsey MG, Bartlett A, Nery JR, Galli M, Gallavotti A, Ecker JR: Cistrome and Epicistrome Features Shape the Regulatory DNA Landscape. Cell 2016, 165(5):1280–1292.

5. Brandt R, Salla-Martret M, Bou-Torrent J, Musielak T, Stahl M, Lanz C, Ott F, Schmid M, Greb T, Schwarz M et al: Genome-wide binding-site analysis of REVOLUTA reveals a link between leaf patterning and light-mediated growth responses. The Plant journal: for cell and molecular biology 2012, 72(1):31–42.

